# The *Bordetella* effector protein BteA induces host cell death by disruption of calcium homeostasis

**DOI:** 10.1101/2024.07.17.603939

**Authors:** Martin Zmuda, Eliska Sedlackova, Barbora Pravdova, Monika Cizkova, Ondrej Cerny, Tania Romero Allsop, Tomas Grousl, Ivana Malcova, Jana Kamanova

## Abstract

*Bordetella pertussis* is the causative agent of whooping cough in humans, a disease that has recently experienced a resurgence. In contrast, *Bordetella bronchiseptica* infects the respiratory tract of various mammalian species, causing a range of symptoms from asymptomatic chronic carriage to acute illness. Both pathogens utilize type III secretion system (T3SS) to deliver the effector protein BteA into host cells. Once injected, BteA triggers a cascade of events leading to caspase 1-independent necrosis through a mechanism that remains incompletely understood. We demonstrate that BteA-induced cell death is characterized by the fragmentation of the cellular endoplasmic reticulum and mitochondria, the formation of necrotic balloon-like protrusions, and plasma membrane permeabilization. Importantly, genome-wide CRISPR-Cas9 screen targeting 19,050 genes failed to identify any host factors required for BteA cytotoxicity, suggesting that BteA does not require a single nonessential host factor for its cytotoxicity. We further reveal that BteA triggers rapid and sustained influx of calcium ions, which is associated with organelle fragmentation and plasma membrane permeabilization. The sustained elevation of cytosolic Ca^2+^ levels results in mitochondrial calcium overload, mitochondrial swelling, cristolysis, and loss of mitochondrial membrane potential. Inhibition of calcium channels with 2-APB delays both the Ca^2+^ influx and BteA-induced cell death. Our findings indicate that BteA exploits essential host processes and/or redundant pathways to disrupt calcium homeostasis and mitochondrial function, ultimately leading to host cell death.

**Importance:** The respiratory pathogens, *Bordetella pertussis* and *Bordetella bronchiseptica*, exhibit cytotoxicity towards a variety of mammalian cells, which depends on the type III secretion effector BteA. Moreover, the increased virulence of *B. bronchiseptica* is associated with enhanced expression of T3SS and BteA effector. However, the molecular mechanism underlying BteA cytotoxicity is elusive. In this study, we performed a CRISPR-Cas9 screen, revealing that BteA-induced cell death depends on essential or redundant host processes. Additionally, we demonstrate that BteA disrupts calcium homeostasis, which leads to mitochondrial dysfunction and cell death. These findings contribute to closing the gap in our understanding of the signaling cascades targeted by BteA.

## Introduction

*Bordetella pertussi*s and *Bordetella bronchiseptica,* known as classical bordetellae, colonize the ciliated epithelium of the respiratory tract of various mammals, inducing a spectrum of respiratory symptoms (1, 2). *B. pertussis*, adapted strictly to humans, is the causative agent of pertussis or whooping cough, which was notorious for its high mortality rates in infants before the introduction of the whole cell pertussis vaccine (wP). The implementation of the wP vaccine substantially reduced pertussis cases. However, with the switch from wP to the acellular pertussis vaccine (aP), there has been a resurgence of pertussis, particularly in adolescents (3). Factors likely contributing to this resurgence include waning immunity in aP-vaccinated individuals and the failure of aP vaccines to elicit Th1/Th17 responses, which is crucial for preventing *B. pertussis* infection, carriage, and transmission (4–6). In contrast, *B. bronchiseptica* has a broader host range but rarely infects humans. Infections caused by *B. bronchiseptica* can range from asymptomatic chronic respiratory carriage to acute illnesses. In dogs, *B. bronchiseptica* causes kennel cough, which is characterized by tracheobronchitis that can progress to bronchopneumonia, especially in puppies (7, 8). In pigs, the bacterium has been associated with atrophic rhinitis, tracheitis, bronchitis, and pneumonia (1, 2, 9).

Cell death is a critical component of host defense against infection, as it eliminates infected cells and triggers immune responses. However, bacterial pathogens have developed sophisticated mechanisms to exploit cell death for their own benefit, facilitating immune evasion and/or nutrient acquisition. The occurrence and manner of cell death are influenced by various factors such as the stage of infection, intensity, host cell type, and physiological state (10, 11). Numerous virulence factors of *B. pertussis* and *B. bronchiseptica* have been identified as inducers of cellular cytotoxicity in a variety of host cells and contribute significantly to the pathogenicity of these bacteria (12–16). One such virulence factor is the type III secretion system (T3SS), which enables the delivery of the cytotoxic effector protein BteA directly from bacterial cytosol into host cells (17, 18). The activity of the T3SS in *B. bronchiseptica* is crucial for persistent colonization of the lower respiratory tract in rats, mice, and pigs, presumably mediated by the actions of the BteA effector (19–21). In addition, increased expression of T3SS genes was associated with the enhanced virulence of the complex I *B. bronchiseptica* 1289 strain isolated from a diseased host compared to the RB50 strain isolated from an asymptomatic host (22, 23). Furthermore, the BteA effector mediates the hypervirulence of complex IV *B. bronchiseptica* isolates (24). Interestingly, the cytotoxicity of BteA of *B. pertussis* is attenuated compared to the considerable BteA-mediated cytotoxicity of *B. bronchiseptica* due to the insertion of an additional alanine at position 503, which may represent an evolutionary adaptation of *B. pertussis* (25). Nevertheless, the role of BteA and T3SS activity in the pathophysiology of pertussis in humans is not yet fully understood.

The 69-kDa effector protein BteA of *B. pertussis* and *B. bronchiseptica* has a modular architecture, consisting of two functional domains: an N-terminal localization/lipid raft targeting (LRT) domain of approximately 130 amino acid residues and a cytotoxic C-terminal domain of approximately 526 and 528 amino acid residues, respectively (26). The N-terminal LRT domain binds negatively charged membrane phospholipids and has a tertiary fold reminiscent of the plasma membrane-targeted 4HBM domain found in a variety of bacterial toxins, including clostridial glucosyltransferase toxins and multifunctional-autoprocessing RTX toxins (MARTX) (26–29). In contrast, the C-terminal cytotoxic domain lacks structural homologs and is solely responsible for cytotoxicity. Indeed, ectopically expressed BteA remains cytotoxic even upon deletion of the LRT domain or the removal of the first 200 N-terminal amino acids (26, 30). The cell death induced by BteA is characterized by its rapid onset, non-apoptotic nature, and independence from caspase-1 (16), although the underlying processes and mechanism of BteA action are unknown. In this work, we demonstrate that BteA induces host cell death by disruption of calcium homeostasis while exploiting essential processes and/or redundant signaling pathways for its necrotic effects.

## Results

### BteA-induced cell death is characterized by rapid fragmentation of the endoplasmatic reticulum and mitochondrial networks

To elucidate the mechanisms underlying BteA-induced host cell death, we first examined the morphological changes in epithelial HeLa cells during infection with *B. bronchiseptica* RB50 strain at multiplicity of infection (MOI) of 10:1. Using time-lapse imaging, we monitored the changes in HeLa cells upon contact with wild-type *B. bronchiseptica* (*Bb*WT) expressing the fluorescent protein mNeonGreen. Shortly after contact, we observed granulation in the cytoplasm of HeLa cells, followed by the formation of plasma membrane blebs, as depicted in Fig. 1A. These blebs continued to expand without retracting, reaching diameters of tens of µm. At the same time, we observed shrinkage of the nucleus and its staining with propidium iodide, indicating early damage to the plasma membrane (Fig. 1A, Suppl. Video 1). In contrast, HeLa cells infected with a *B. bronchiseptica* derivative lacking the *bteA* gene (*Bb*Δ*bteA*) and expressing mNeonGreen at the same MOI exhibited no such morphological changes (Fig. 1A, Suppl. Video 1).

**Figure 1.**
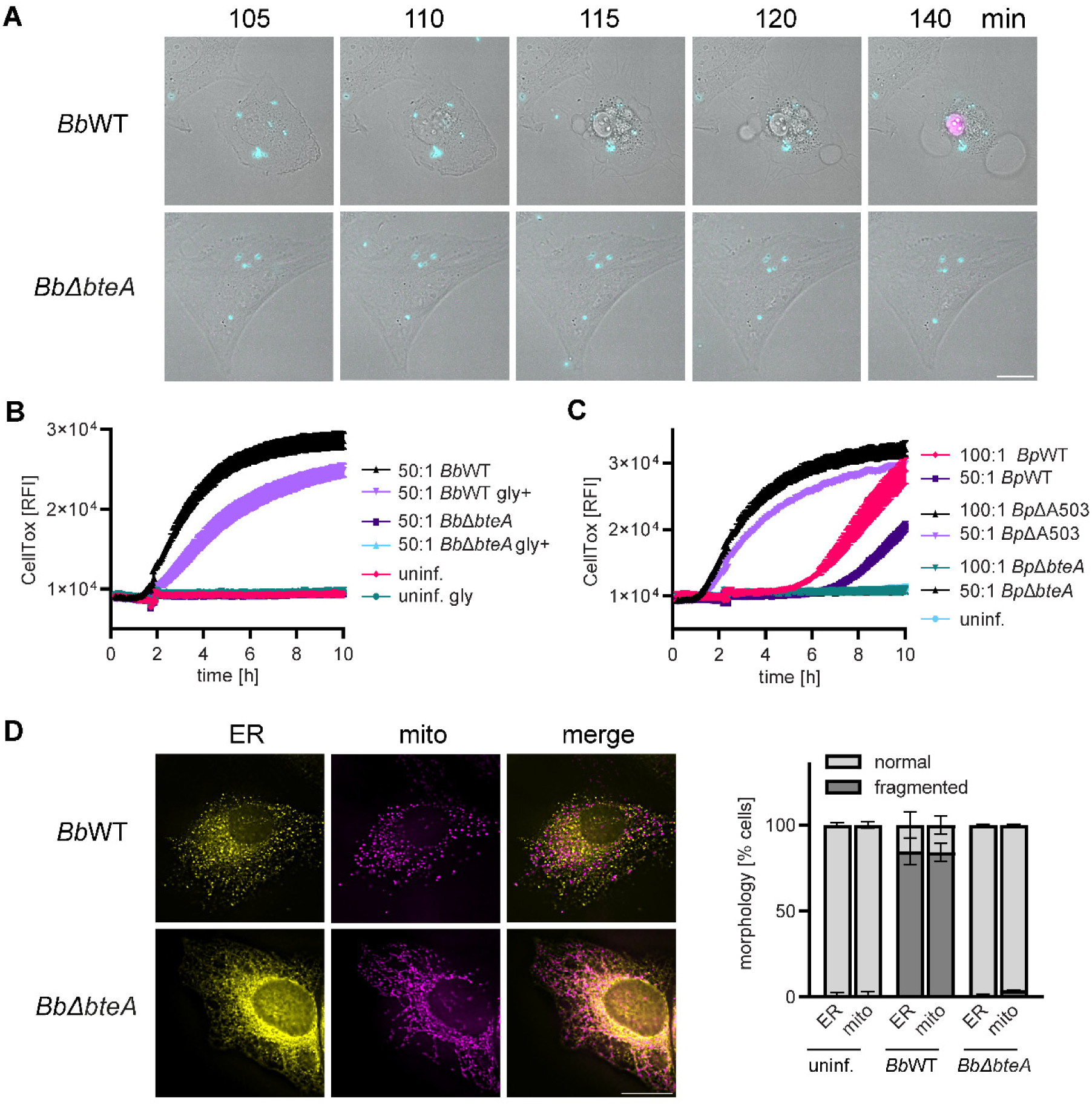
BteA-induced cell death is characterized by fragmentation of endoplasmatic reticulum and mitochondrial networks. **(A)** Time lapse analysis of morphological changes and plasma membrane permeabilization in Hela cells. Hela cells were infected with *B. bronchiseptica* wild type (*Bb*WT) and *Bb*Δ*bteA* mutant (*Bb*Δ*bteA*), expressing the fluorescent protein mNeonGreen, at MOI of 10:1 in the presence of propidium iodide (5 µg/ml). Sequence of time lapse images is shown. Data are representative of three independent experiments. Bright field, gray; bacteria, cyan; propidium iodide, magenta. Scale bar, 20 µm. **(B-C)** Comparison of *B. bronchiseptica* and *B. pertussis* cytotoxicity against HeLa cells. HeLa cells were infected with *B.bronchiseptica* **(B)**, or *B. pertussis* **(C)** wild type strains, and their mutant derivatives at the indicated MOI. Plasma membrane permeabilization was determined using the fluorescent DNA binding dye CellTox Green. For *B. bronchiseptica*, infections were conducted both in the presence (gly+) and absence of glycine (5 mM). Data represent the mean ± SEM of a representative experiment from 2 independent experiments. **(D)** Visualization of cellular structures. Hela cells were transfected to express fluorescent proteins tagged with localization signals for endoplasmatic reticulum (ER) and mitochondria (mito). One hour after infection with *Bb*WT and *Bb*Δ*bteA* mutant at MOI 50:1, cells were fixed and analyzed by fluorescence imaging. The shown micrographs are representative of two independent experiments from which the organelle morphology was determined. Data represent the mean ± SEM of at least 100 cells counted per experiment. ER, yellow; mitrochondria, magenta. Scale bar, 20 µm.

We confirmed these observations by following the kinetics of plasma membrane permeabilization using the fluorescent DNA binding dye CellTox Green. Infection of HeLa cells with *Bb*WT bacteria at MOI of 50:1 resulted in disruption of cytoplasmic membrane integrity, whereas the *Bb*Δ*bteA* mutant strain failed to induce membrane permeabilization, showing that the observed cytotoxicity of the RB50 strain against HeLa cells was due to the action of the BteA effector (Fig. 1B). As previously reported and also shown in Fig. 1B, we could delay BteA-induced membrane permeabilization by glycine, a plasma membrane-stabilizing amino acid that blocks the opening of non-specific anion channels that lead to colloid osmotic swelling and plasma membrane failure (16). Wild type *B. pertussis* B1917 (*Bp*WT) at MOI of 50:1 also induced BteA-dependent plasma membrane permeabilization of HeLa cells, but this occurred 6 hours post-infection compared to 2 hours with *Bb*WT (Fig. 1C). Moreover, increasing the *Bp*WT MOI to 100:1 accelerated this onset by only 2 hours. Importantly, as previously demonstrated (25), removing A503 from the BteA effector of *B. pertussis* B1917 (*Bp*ΔA503) resulted in a loss of plasma membrane integrity comparable to that of *Bb*WT (Fig. 1C). These results confirmed the activity of T3SS in *B. pertussis* and the critical role of A503 in BteA cytotoxicity.

To corroborate the processes occurring in HeLa cells just before the BteA-induced plasma membrane blebbing and permeabilization, we used plasmid-encoded fluorescent markers targeted to the endoplasmic reticulum (ER) and mitochondria, specifically export&KDEL-mScarlet and 4xmts-mNeonGreen (31). HeLa cells were transiently transfected with these markers, and infected with either *Bb*WT and *Bp*WT strains or their derivatives at MOI 50:1. Cells were fixed at 1 h or 5 h post-infection for fluorescence microscopy analysis. As shown in Fig. 1D, extensive fragmentation and vesiculation of the ER and mitochondrial networks was observed in *Bb*WT-infected cells compared to those infected with *Bb*Δ*bteA* derivative or uninfected cells as early as 1 h post-infection. Within 1 h of infection, *Bp*ΔA503 induced similar fragmentation of the ER and mitochondria as *Bb*WT while no extensive fragmentation was detected in Hela cells infected by *Bp*WT (Suppl. Fig. 1 A). However, the fragmentation induced by *Bp*WT compared to controls became pronounced at 5 hours post-infection, just before plasma membrane permeabilization and cell death, as further shown in Suppl. Fig. 1B.

In summary, our data demonstrate that BteA from both *B. bronchiseptica* and *B. pertussis* induces fragmentation of ER and mitochondrial networks, which precedes colloid osmotic lysis of HeLa cells. The induction of ER and mitochondrial fragmentation and cell death occurred faster during infection with *B. bronchiseptica* due to the absence of A503 in the BteA effector.

### BteA targets essential components of the host cell or genes with redundant functions

To identify host factors essential for BteA-induced cytotoxicity and potentially elucidate the signaling cascades responsible for the observed fragmentation of ER and mitochondria, we next performed a CRISPR-Cas9 forward genetic screen (32–34). This screen employed human embryonic kidney HEK cells, which are similarly sensitive to BteA action as HeLa cells (Suppl. Fig. 2A) but are more suitable for this approach. To generate Hek-Cas9 mutant library, we used commercially available GeCKO v2 library (35), divided into sublibraries A and B, each targeting approximately 19,000 human genes with three distinct sgRNAs. We packaged these sgRNAs into lentiviral particles and introduced them into Cas9-expressing HEK (HEK-Cas9) cells with MOI of 0.3 to ensure the delivery of a single sgRNA per cell, as depicted in Fig. 2A. Subsequent sequencing of generated Hek-Cas9 sublibraries A and B showed a compact sgRNA distribution, containing the vast majority of sgRNAs in hundreds of hits to the 15 million normalized reads (Suppl. Fig. 2 B and C). When comparing the number of targeted genes in the respective sublibraries to the theoretically present number of targeted genes, we found that 54 genes in sublibrary A and 53 genes in sublibrary B were not targeted (Suppl. Fig. 2 B and C). However, due to overlap in targeted genes, only 10 genes were not targeted in the combined sublibraries A and B, as shown Fig. 2B. These genes, listed in Suppl. Table S1, include the phospholamban gene, 8 miRNAs, and 1 non-targeting control guide. Furthermore, as shown in the comparison between Fig. 2B and Suppl. Fig. 2D, if a gRNA was not detected, it typically constituted one of the six gRNAs present in the theoretical pool. Overall, our analysis of the resulting HEK-Cas9 sublibraries A and B confirmed the successful introduction of sgRNAs of GeCKO v2 library and preservation of library complexity.

**Figure 2.**
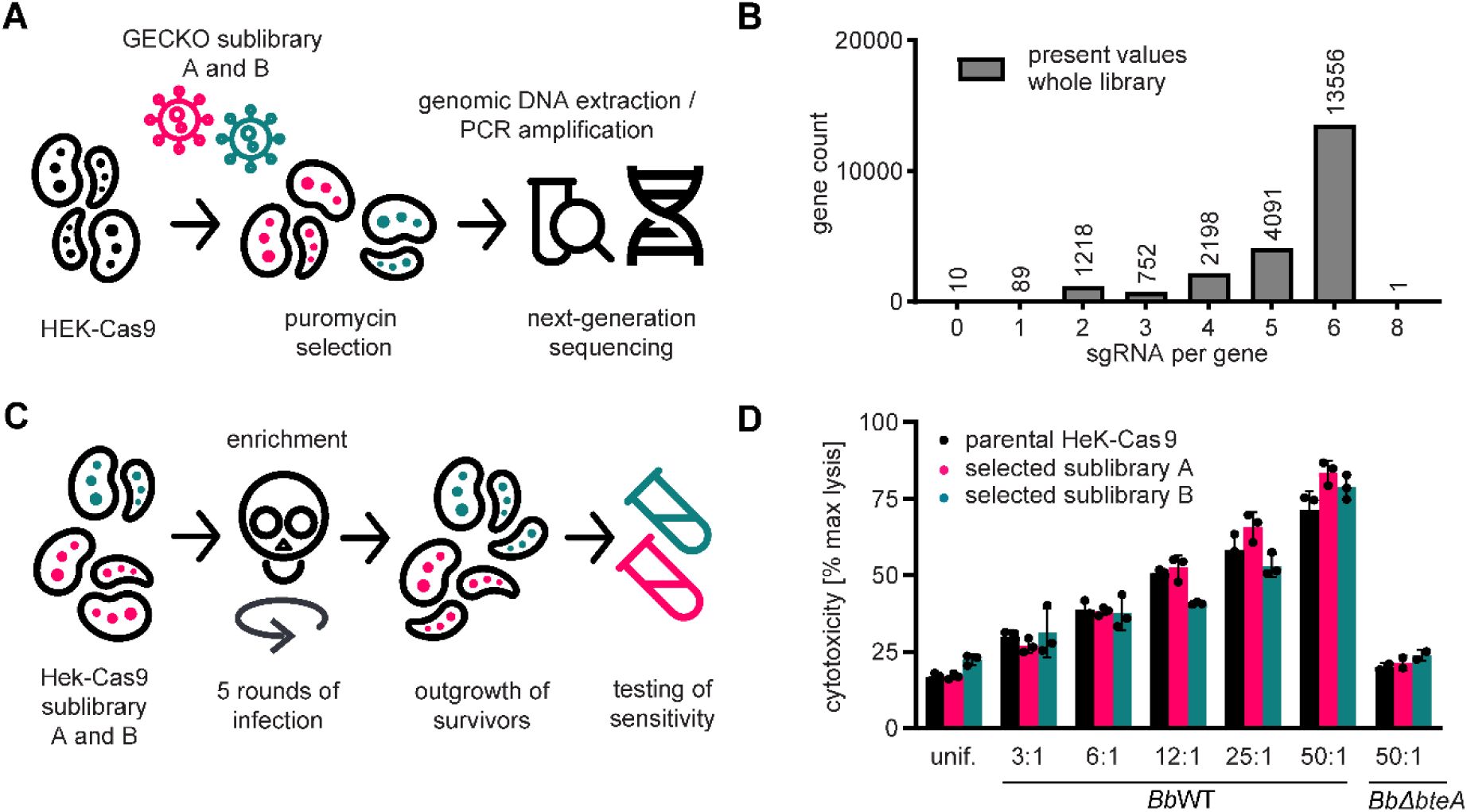
BteA targets essential components of the host cell or genes with redundant functions. **(A)** Workflow for generating the Hek-Cas9 sublibraries A and B. Sublibraries A and B of the GeCKO v2 library were individually packaged into lentiviral particles and introduced into Cas9-expressing HEK cells. After selection with puromycin and expansion of both cell pools, genomic DNA was extracted, sgRNA amplified and barcoded, and library complexity was verified by next-generation sequencing. **(B)** Verification of library complexity in the Hek-Cas9 sublibraries A and B. The number of targeted genes per detected sgRNA count is indicated. **(C)** Workflow for enrichment of BteA-resistant cells. Hek-Cas9 sublibraries A and B were subjected to five rounds of selection with *Bb*WT at MOI 100:1 for 5 h per round. Each selection resulted in the death of 95% of infected cells. To stop the infection, the medium containing *Bb*WT was discarded and replaced with fresh medium containing 100 µg/ml of gentamicin. In between selection rounds, the surviving HEK-Cas9 sublibraries A and B were expanded. **(D)** Susceptibility of selected Hek-Cas9 sublibraries A and B. Selected Hek-Cas9 sublibraries A and B, and parental Hek-Cas9 cells were infected with *Bb*WT at the indicated MOI. The cytotoxicity was determined as lactate dehydrogenase (LDH) release 6 h post-infection. Data represent the mean ± SEM of a representative experiment from 2 independent experiments.

Subsequently, the Hek-Cas9 cell pools were exposed to *Bb*WT at MOI 100:1 for 5 h before *Bb*WT was eliminated by gentamicin. These conditions were previously determined in pilot experiments, and induced death in 95% of infected cells. The surviving cells were expanded, and the process was repeated for five rounds of infection to enrich for resistant cells, as depicted in Fig. 2C. Our hypothesis was that infection of transduced cells with *Bb*WT would enrich for sgRNA-directed mutations, which lead to resistance against BteA action due to inactivation of genes required for cell death. However, when comparing the cell pools that survived five rounds of *Bb*WT infection with the parental HEK-Cas9 cells (Fig. 2D), no acquired resistance was detected. Thus, our CRISPR-Cas9 knockout screen failed to identify host genes essential for BteA function. Possible explanations include redundancy of signaling pathways and lack of non-essential targets.

### BteA disrupts cytosolic calcium homeostasis, leading to permeabilization of cell plasma membrane

Given that the CRISPR-Cas9 screen did not provide mechanistic insights and previous research has shown that changes in the structure of the ER and mitochondria are linked to increased cytosolic calcium levels (36, 37), we investigated whether BteA disrupts cellular calcium homeostasis. To this end, HeLa cells were loaded with the single-wavelength fluorescent Ca^2+^ indicator, Fluo-4/AM (38) and subjected to fluorescence-intensity measurement at 516 nm after infection with *B. bronchiseptica*. As shown in Fig. 3A, *Bb*WT induced cytosolic Ca^2+^ influx, with the intensity and onset of this influx varying based on the used MOI. At MOIs of 50:1 and 25:1, the Ca^2+^ influx peaked around 1 h post-infection, which was followed by a decrease in signal, whereas at MOI of 5:1, the peak of Ca^2+^ influx occurred approximately 2 hours after infection. At each MOI, the peak of Ca^2+^ influx induced by *Bb*WT infection preceded the permeabilization of HeLa cell plasma membrane, as shown by the binding of the DNA dye CellTox Green (Fig. 3B). Importantly, no such Ca^2+^ influx was observed when HeLa cells were infected with the *Bb*Δ*bteA* derivative at the same MOIs, confirming the essential role of BteA in this process.

**Figure 3.**
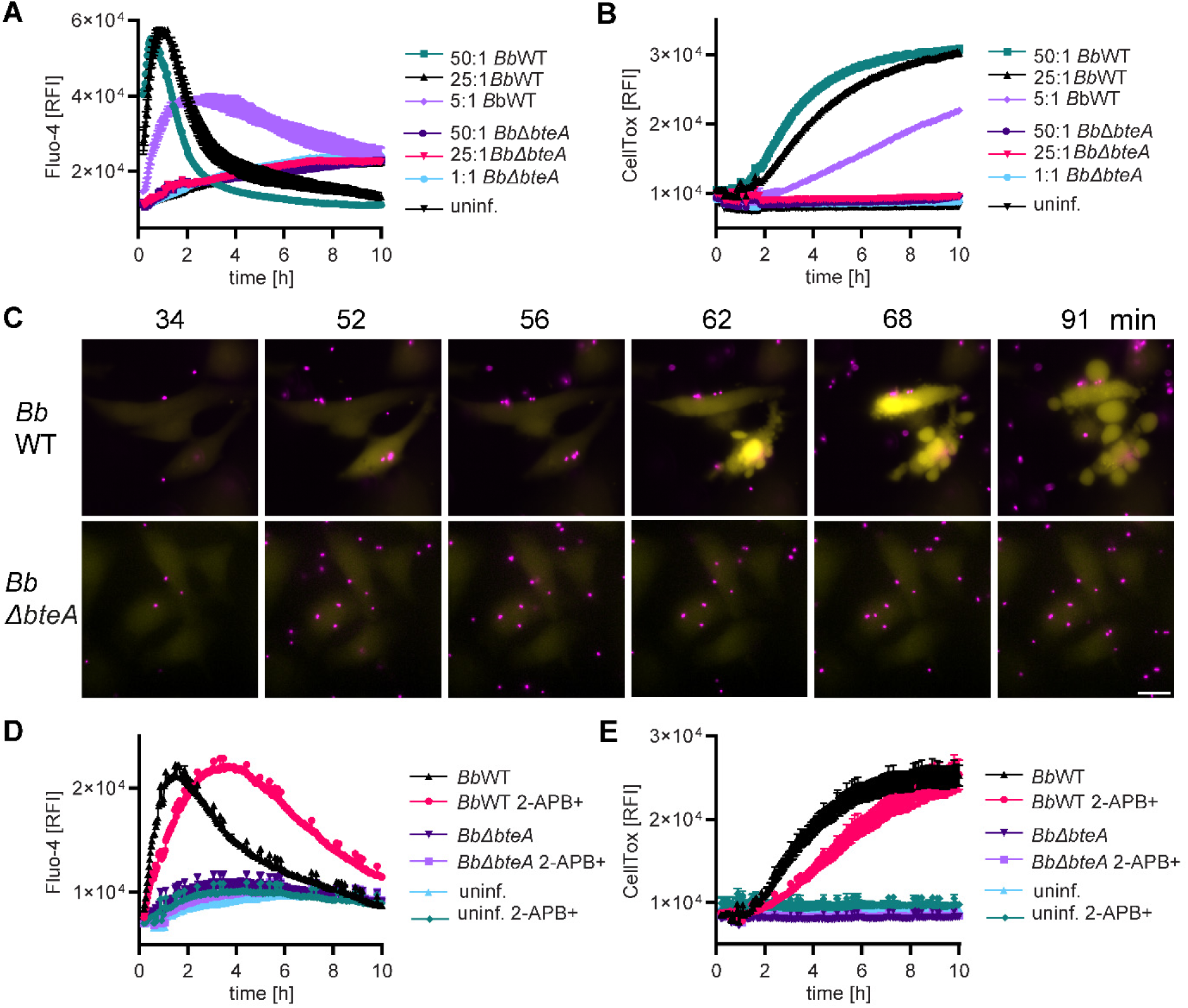
BteA disrupts cytosolic calcium homeostasis, leading to permeabilization of cell plasma membrane. **(A-B)** Calcium influx precedes *B. bronchiseptica* induced-plasma membrane permeabilization. HeLa cells were infected with *Bb*WT and *Bb*Δ*bteA* mutant at the indicated MOI. Calcium influx was monitored using cytosolic Ca^2+^ indicator Fluo-4/AM **(A)**, while plasma membrane permeabilization was determined in parallel wells by fluorescent DNA binding dye CellTox Green **(B)**. Data represent the mean ± SEM of a representative experiment from 3 independent experiments. **(C)** Calcium imaging. HeLa cells loaded with cytosolic Ca^2+^ indicator Fluo-4/AM were infected with *Bb*WT and *BbΔbteA*, expressing the fluorescent protein mScarlet (mSc), at MOI of 10:1. Sequence of time lapse images is shown. Data are representative of two independent experiments. Bacteria, magenta; cytosolic Ca^2+^ indicator Fluo-4/AM, yellow. Scale bar, 20 µm. **(D-E)** Treatment with 2-APB delays calcium influx and host cell death. HeLa cells were pre-incubated with 100 µM 2-APB (2-APB+) for 30 min, or left untreated, before being infected with *Bb*WT or *Bb*Δ*bteA* derivative at MOI 25:1. Calcium influx was in assessed using Fluo-4/AM Ca^2+^ indicator **(D)** whereas plasma membrane permeabilization was determined in parallel wells by fluorescent DNA binding dye CellTox Green **(E)**. Data represent the mean ± SEM of a representative experiment from 3 independent experiments.

Subsequently, the response of individual cells to infection with *Bb* strains expressing mScarlet (mSc) at MOI 10:1 was visualized through time-lapse imaging of Fluo-4/AM-loaded HeLa cells. As shown in Fig. 3C and Suppl. Video 2, the attachment of *Bb*WT to the HeLa cell surface triggered a sustained increase in cytosolic Ca^2+^ levels, which was detected as an increase in Fluo-4/AM fluorescence intensity as early as 30 min after contact. This was followed by BteA-induced plasma membrane blebbing, permeabilization and cell death. The loss of membrane integrity was noticeable as a decrease in Fluo-4/AM fluorescence intensity. Indeed, inhibition of plasma membrane permeabilization by glycine did not affect the onset of Ca^2+^ influx but prolonged the duration of the Fluo-4/AM signal, as shown in Suppl. Fig. 3 A and B. In a subset of cells, *Bb*WT infection induced cytosolic Ca^2+^ oscillations, as shown in Suppl. Video 3, which did not immediately lead to the plasma membrane blebbing being strictly associated with sustained increase in cytosolic calcium levels. Importantly, no major changes in cytosolic Ca^2+^ were observed after infection with *Bb*Δ*bteA* at the same MOI (Fig. 3C, Suppl. Video 2 and 3).

To determine the contribution of disrupted cellular calcium homeostasis to BteA-induced cell death, we next analyzed whether the commonly used and rather unspecific calcium channel modular, 2-aminoethyl diphenylborinate (2-APB), could block BteA-induced cytosolic Ca^2+^ influx and cell death. Importantly, pretreatment with 100 μM 2-APB for 30 min delayed the cytosolic Ca^2+^ influx mediated by BteA, as shown in Fig. 3D. This delay was accompanied by a postponement of BteA-induced membrane permeabilization (Fig. 3E), showing the critical role of cytosolic Ca^2+^ increase in this process.

Overall, our data demonstrate that BteA induces sustained cytosolic Ca^2+^ influx, which triggers cytoplasmic plasma membrane blebbing and permeabilization, ultimately leading to host cell death.

### BteA-induced calcium influx is associated with ER and mitochondria fragmentation and yields elevation of mitochondrial calcium levels

To gain insight into BteA-induced Ca^2+^ fluxes, we next utilized cytosolic Fluo-4/AM calcium indicator in combination with plasmids encoding red fluorescent Ca^2+^ sensors targeted either to ER or mitochondria, ER-LAR-Geco and mito-LAR-Geco (39), respectively.

First, HeLa cells were transiently transfected with ER-LAR-Geco, loaded with Fluo-4/AM and subjected to infection with *Bb*WT. The increase in cytosolic Ca^2+^ concentration correlated with fragmentation of ER network, as shown in Fig. 4A and Suppl. Video 4. However, quantification of ER-LAR-Geco fluorescence intensity revealed no substantial changes (Fig. 4A and Suppl. Fig. 4). In contrast, cells transfected to express mito-LAR-Geco (Fig. 4B and Suppl. Fig. 5) showed a simultaneous increase in mitochondrial and cytosolic Ca^2+^ levels. This increase in mitochondrial Ca^2+^ levels preceded mitochondrial fragmentation and plasma membrane blebbing, as evidenced in Fig. 4B and Suppl. Video 5. Consistent results were obtained when cell mitochondria were visualized using MitoTracker (Suppl. Fig. 6 and Suppl. Video 6). Thus, BteA-induced Ca^2+^ influx is associated with the fragmentation of the ER and mitochondria and leads to elevated mitochondrial Ca^2+^ levels.

**Figure 4.**
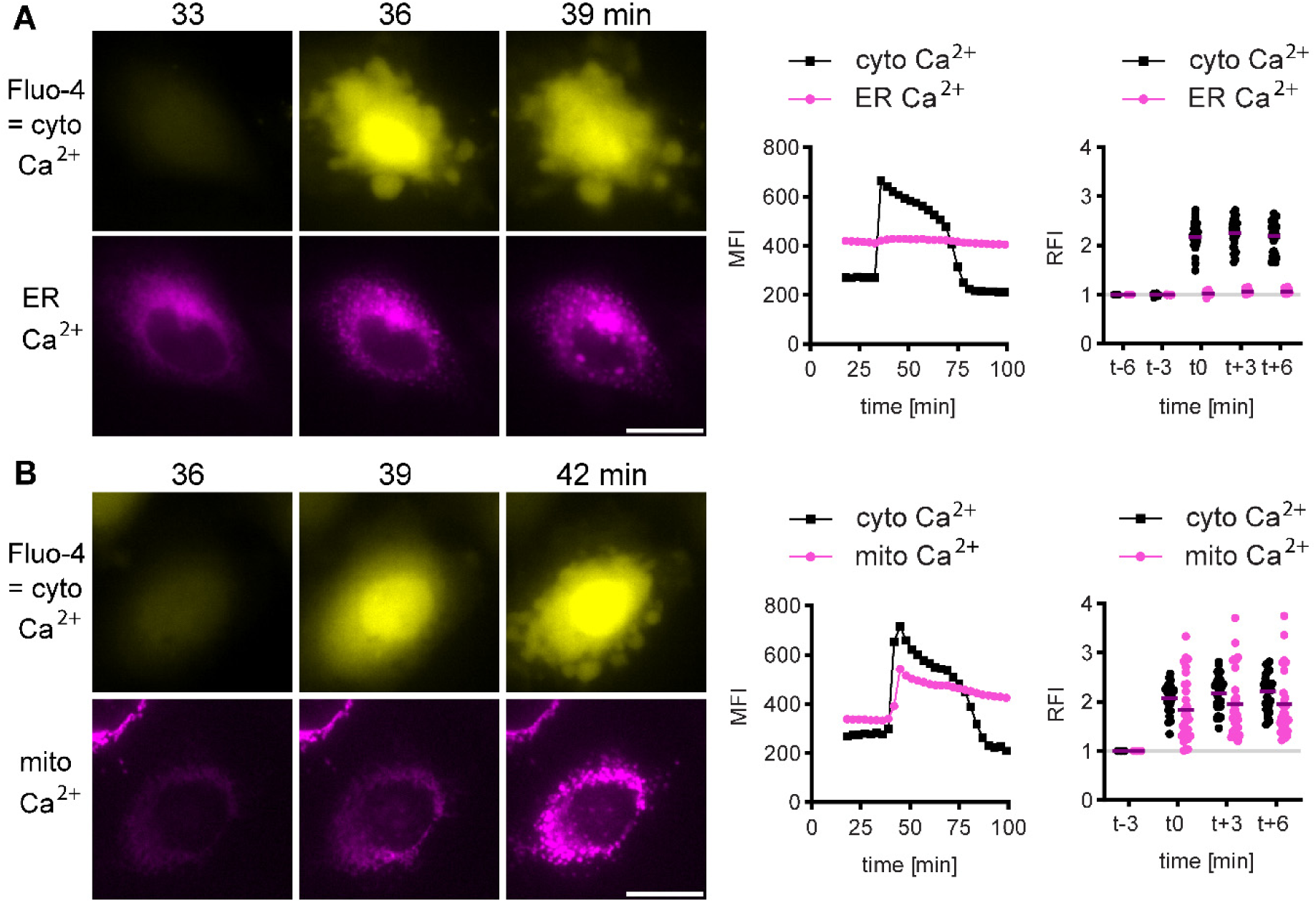
BteA-induced calcium influx correlates with ER and mitochondria fragmentation and yields elevation of mitochondrial calcium levels. **(A)** ER calcium imaging. Hela cells, transfected to express ER-targeted red Ca^2+^ sensor ER-LAR-GECO, were loaded with the cytosolic Ca²⁺ indicator Fluo-4/AM, and infected with *Bb*WT at MOI of 10:1. A sequence of time-lapse images is shown. The graph in the middle depicts the mean fluorescence intensity of the shown cell quantified over time. The right graph shows the relative fluorescence intensities of individual cells (n=30) at the time of calcium influx. Data are representative of two independent experiments. Cytosolic Ca^2+^ indicator Fluo-4/AM, yellow; ER Ca^2+^ sensor ER-LAR-Geco, magenta. Scale bar, 20 µm. **(B)** Mitochondrial calcium imaging. Hela cells, transfected to express mitochondria-targeted red Ca^2+^ sensor mito-LAR-GECO, were loaded with the cytosolic Ca²⁺ indicator Fluo-4/AM, and infected with *Bb*WT at MOI of 10:1. A sequence of time-lapse images is shown. The graph in the middle depicts the mean fluorescence intensity of the shown cell quantified over time. The right graph shows the relative fluorescence intensities of individual cells (n=30) at the time of calcium influx. Data are representative of two independent experiments. Cytosolic Ca^2+^ indicator Fluo-4/AM, yellow; mitochondrial Ca^2+^ sensor mito-LAR-Geco, magenta. Scale bar, 20 µm.

### BteA induces loss of mitochondrial membrane potential which is accompanied by mitochondrial swelling and cristolysis

To test the impact of BteA-induced mitochondrial Ca^2+^ uptake on mitochondrial function, we next evaluated mitochondrial membrane potential using tetramethylrhodamine (TMRM), which accumulates in healthy mitochondria. A loss of mitochondrial membrane potential leads to disappearance of the TMRM signal. Indeed, mitochondria play a critical role in maintaining Ca^2+^ homeostasis by sequestering calcium ions. However, excessive Ca^2+^ uptake by mitochondria, known as mitochondrial calcium overload, can trigger mitochondrial swelling, opening of the mitochondrial permeability transition pore (mPTP) and loss of mitochondrial membrane potential, ultimately leading to cell death (40, 41).

Time-lapse imaging revealed that BteA-induced Ca^2+^ influx leads to a loss of mitochondrial membrane potential (Fig. 5A, Suppl. Figure 7 and Suppl. Video 7). In addition, electron microscopy showed mitochondrial swelling and disappearance of mitochondrial cristae in *Bb*WT-infected cells (Fig. 5B and Suppl. Figure 8). However, we were unable to inhibit BteA-induced plasma membrane permeabilization by ruthenium 360, an inhibitor of the mitochondrial calcium uniporter channel or cyclosporine A, an inhibitor of mPTP (Fig. 5C). These data indicate that BteA-induced mitochondrial Ca^2+^ uptake correlates with mitochondrial failure, but mitochondrial damage is not solely responsible for execution of BteA-induced cell death. Alternatively, treatments aimed at preventing mitochondrial damage might not be effective in counteracting the sustained Ca^2+^ influx caused by BteA. In fact, pre-treatment of cells with ruthenium 360 was not able to prevent loss of mitochondrial membrane potential, as shown in Suppl. Video 8.

**Figure 5.**
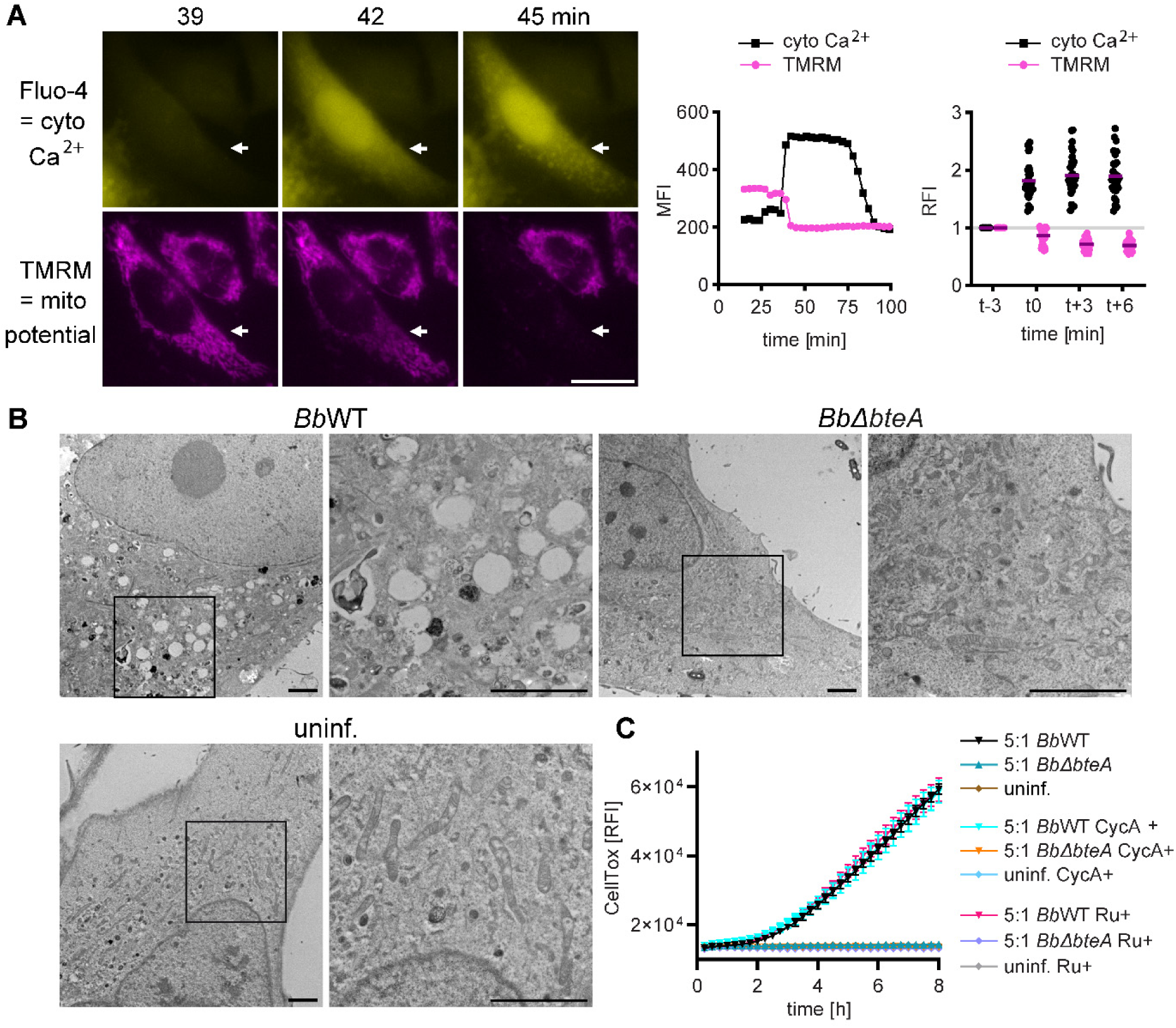
BteA induces loss of mitochondrial membrane potential, mitochondrial swelling and cristolysis. **(A)** BteA-induced calcium influx precedes loss of mitochondrial membrane potential. HeLa cells were loaded with the mitochondrial membrane potential indicator TMRM and the cytosolic Ca*^2+^* indicator Fluo-4/AM, which was followed by infection with *Bb*WT at MOI 10:1. Sequence of time lapse images is shown. The graph in the middle depicts the mean fluorescence intensity of the shown cell (white arrow) quantified over time. The right graph shows the relative fluorescence intensities of individual cells (n=30) at the time of calcium influx. Data are representative of two independent experiments. Cytosolic Ca^2+^ indicator Fluo-4/AM, yellow; TMRM, magenta. Scale bar, 20 µm. **(B)** Transmission electron microscopy images of mitochondria. Hela cells were infected with either *B*bWT or *Bb*Δ*bteA* at MOI of 25:1 for 1 h or left untreated. Following fixation, ultrathin sections were prepared and analyzed. The black square highlights the enlarged section. Images are representative of two independent experiments. Scale bar, 2 µm. **(C)** Role of mitochondria failure in the execution of BteA-induced cell death. HeLa cells were pre-incubated with either 1 µM cyclosporin A (CycA+) or 10 µM ruthenium 360 (Ru+) for 1 h, or left untreated, before being infected with *Bb*WT and Δ*bteA* derivative at an MOI 5:1. Plasma membrane permeabilization was determined using the fluorescent DNA binding dye CellTox Green. Data represent the mean ± SEM of a representative experiment from 2 independent experiments.

## Discussion

Our study demonstrates that injection of *Bordetella* type III effector protein BteA into the host cell triggers a cytosolic Ca^2+^ influx that disrupts cellular calcium homeostasis. This sustained increase in cytosolic Ca^2+^ is detectable within minutes of bacterial contact and leads to mitochondrial Ca^2+^ overload, plasma membrane blebbing, and early membrane damage. Furthermore, CRISPR-Cas9 screen shows that BteA targets critical and/or redundant cellular survival mechanisms and does not dependent on a single non-essential host factor for its cytotoxic effects.

The type III secretion effector BteA of classical *B. bronchiseptica* and *B. pertussis* is a potent cytotoxin that causes cell death in a variety of cells, including epithelial cells and macrophages (16, 17, 19, 42). Previous studies have shown that BteA induces morphological changes in rat epithelial lung L2 cells within just 25 minutes (42, 43). In our time-lapse experiments, when we centrifuged *B. bronchiseptica* at MOI of 10:1 onto HeLa cells and examined the cells, we detected increased levels of cellular Ca^2+^ already during the initial microscope focusing. No more than 10 minutes after adding the bacteria to the host cells. Similarly, spectrophotometric measurements of a whole cell population infected with MOI of 25:1 showed elevated levels of Ca^2+^ at the first collected data point. When the bacteria during time-lapse microscopy were added without centrifugation so that we could follow their sedimentation on the cell surface, we observed a sustained increase in cytosolic Ca^2+^ as early as 30 minutes after bacterial contact with the host cell. The plasma membrane blebbing and permeabilization were observed after another 30 to 60 minutes. These results demonstrate the efficiency of T3SS-mediated BteA delivery into the host cell and BteA potency to disrupt cellular signaling and cause cell death.

The most prominent feature of the BteA-induced morphological changes during cell death was the swelling of the cytoplasm and the formation of balloon-like, non-retracting plasma membrane blebs. This observation is consistent with previous studies (16, 30) and is a common manifestation of cell injury associated with a lytic form of cell death. In fact, regulated cell death can be categorized into lytic and non-lytic forms. Lytic forms such as pyroptosis and necroptosis, involve the formation of pores or breaches in the plasma membrane and the expansion of the cytoplasmic compartment due to an ionic imbalance. This process ultimately leads to cell rupture and the release of intracellular contents into the extracellular space. In contrast, non-lytic forms, such as apoptosis, involve the coordinated disintegration of dying cells into smaller fragments without compromising the integrity of the cytoplasmic membrane (44). Similar to necrosis, apoptosis is also characterized by blebbing of the plasma membrane. In apoptotic blebbing, however, there are repeated protrusions and retractions of the plasma membrane, which lead to the formation of apoptotic bodies. In necrotic blebbing, on the other hand, no retractions take place, allowing blebs to grow to larger (45). The signaling processes underlying blebbing are complex. Key events in the formation of blebs include disruption of the actomyosin cell cortex and its interaction with the plasma membrane, and regulation of cytoplasmic fluidity. Importantly, Ca^2+^ is important regulator of actin-binding proteins that control the dynamics of the actin cytoskeleton and are involved in the regulation of bleb formation (46, 47). An increase in Ca^2+^ leads to the fragmentation of F-actin through the activation of gelsolin and INF2 (48, 49) and also increases cytoplasmic fluidity in the expanding bleb (50). In addition, Ca^2+^ activate calpains, cytoplasmic cysteine proteases of the papain family, which cleave various substrates, including cytoskeletal proteins such as spectrin, E-cadherin and vinculin, and thus contribute to cell damage (51, 52).

It is important to point out that a sustained increase in cytosolic Ca^2+^ is an upstream event in BteA-induced cytotoxicity. First, this increase in cytosolic Ca^2+^ happens quickly and precedes any nonspecific damage to the plasma membrane. Second, inhibition of plasma membrane permeabilization with glycine, which blocks the opening of nonspecific anion channels, has no effect on the onset of Ca^2+^ influx. Finally, the calcium channel modulator 2-APB delays both Ca^2+^ influx and plasma membrane permeabilization. Interestingly, time-lapse experiments with fluorescence images taken every 3 minutes also showed that infection with *Bb*WT, in contrast to infection with the mutant *Bb*Δ*bteA* strain, induces cytoplasmic Ca^2+^ oscillations in a subset of HeLa cells. These oscillations occurred both in the cells prior to the sustained Ca^2+^ increase and in the cells that did not show a sustained Ca^2+^ increase during the imaging period. We hypothesize that these oscillations may be a coping mechanism of the cells in response to BteA action. Alternatively, it could be a response to ATP release from neighboring lysed cells, which has been described as a trigger for Ca^2+^ oscillations (53). The relationship of these oscillations to BteA-induced cell death remains to be investigated.

The 69 kDa BteA effector comprises two different functional domains. The N-terminal region, consisting of 130 amino acids, which binds to negatively charged membrane phospholipids and localizes BteA to the plasma membrane (26, 28, 29), and the C-terminal region, consisting of 526 amino acids in *B. pertussis* and 528 amino acids in *B. bronchiseptica*, which is solely responsible for cytotoxicity (25, 26). No known homologs exist for the C-terminal region, and prediction and homology search tools have been unable to elucidate the mechanism underlying its cytotoxicity (18). Even models generated by the AlphaFold prediction server (54) exhibit low confidence in the structural prediction of this region. The kinetics of cell death induced by *Bb*WT and *Bp*WT strains can be linked to the presence or absence of alanine 503 in the BteA effector, although it is unclear how this insertion affects BteA structure (25). Analyses using transmembrane predictors such as DeepTMHMM (55), https://dtu.biolib.com/DeepTMHMM, indicate no transmembrane alpha-helical or beta-barrel segments within the BteA molecule, suggesting it does not form membrane pores. Consequently, potency and rapid effect of BteA on disrupting calcium ion homeostasis suggest that BteA either affects host cell enzymes and/or membrane channels, or possesses enzymatic activity itself.

The delivery of BteA into the host cell cytosol is required for cytotoxicity. Mutant strains of *B. bronchiseptica* lacking translocon components such as BopB, BopD and Bsp22, which are required for the translocation of BteA into the host cells (but not for its secretion into the cell medium), do not induce cell death (43, 56–58). In contrast, heterologous expression of BteA causes cytotoxicity in both mammalian cells and *S. cerevisiae* (25, 26), which indicates that the pathways activated by BteA that lead to cytotoxicity are conserved across species. Our CRISPR-Cas9 forward genetic screen, which applied stringent selective pressure, further suggests that the host cell factors targeted by BteA are essential and/or that the mechanism of BteA action is redundant. This screen aimed to identify genes that are required for the cytotoxicity of BteA by inactivating them using the GeCKO v2 library (35). Despite the successful introduction of the sgRNA library, targeting approximately 19,000 genes, into Hek-Cas9 cells, we were unable to select resistant cells and pinpoint specific host genes essential for BteA-induced cell death. More refined or complementary genetic screens might be necessary to identify host factors, which contribute redundantly and/or partially. These screens should use lower selective pressure to better differentiate between cells with and without the target gene knockouts, although this approach also presents challenges. The amount of BteA delivered by bacteria is not precisely controlled, resulting in variable delivery among cells, and using a stable cell line with inducible BteA expression is also not straightforward. We have been unsuccessful in constructing a stable Hek-Cas9 cell line expressing doxycycline-inducible *Bb*BteA due to promoter leakiness. Additionally, while we managed to construct a Hek-Cas9 line with inducible expression of the less toxic *Bp*BteA, this cell line was unstable, leading to loss of the *BpbteA* gene after several passages.

The lack of identified genes required for BteA cytotoxicity aligns well with the observations that BteA rapidly disrupts calcium homeostasis, which is a crucial cellular signaling mechanism. Indeed, calcium ions serve as a universal cellular messenger but can also become toxic and cause cell death. In a resting cell, the cytoplasmic Ca^2+^ concentration is relatively low, about 100 nM, similar to that in the mitochondria. In contrast, the Ca^2+^ concentration in the ER is between 100 μM and 1 mM, making the ER the main calcium ion reservoir in the cell and a crucial player in coordination with other organelles and the plasma membrane (59, 60). During cell stimulation, the cytosolic Ca^2+^ concentration increases approximately 10 times to 1 μM due to the release of Ca^2+^ from the ER or the influx of extracellular Ca^2+^. In addition, the ER transmits Ca^2+^ signals to the mitochondria via transport systems located at the mitochondria-associated ER membranes (MAMs) and relying on the IP3 receptors (IP3R) and the mitochondrial calcium uniporter (MCU) (61, 62). To gain insight into the disruption of calcium homeostasis by the action of BteA, we monitored cellular Ca^2+^ fluxes triggered by BteA. Although we observed oscillatory cytosolic Ca^2+^ increases, indicating Ca^2+^ release from the ER via IP3R, we did not detect any changes in ER Ca^2+^ concentration. This could be due to two reasons. First, the low-affinity intensiometric red fluorescent Ca^2+^ sensor ER-LAR-Geco might lack the sensitivity to detect changes in ER Ca^2+^ concentration. Second, the 3-minute intervals used to acquire the fluorescence images may have allowed any decrease in ER Ca^2+^ concentration to reverse. Depletion of ER Ca^2+^ stores triggers an influx of Ca^2+^ from the extracellular space, known as capacitative Ca^2+^ influx or store-operated Ca^2+^ entry (SOCE), which increases cytosolic calcium levels and replenishes the ER (59). Future studies should investigate how BteA triggers Ca^2+^ influx and which cellular components are involved. While the calcium channel modulator 2-APB is useful in assessing the general importance of cellular calcium levels, it is not suitable for detailed mechanistic experiments. 2-APB was initially discovered as an inhibitor of the IP3R in the ER but was later also shown to modulate SOCE and block plasma membrane TRPC and TRPM channels (63–66).

Importantly, unlike changes in the ER Ca^2+^ levels, we demonstrate that BteA causes a sustained Ca^2+^ influx into the mitochondria. This influx is crucial as mitochondrial Ca^2+^ dynamics play a critical role in determining cellular outcomes. Rhythmic low-level Ca^2+^ oscillations enhance mitochondrial ATP production, while insufficient Ca^2+^ uptake slows cell proliferation. Furthermore, prolonged Ca^2+^ accumulation can trigger cell death (61). Mitochondria can accumulate 10 to 20 times more Ca^2+^ than the cytosolic compartment, however, excessive Ca^2+^ accumulation can trigger the opening of the mitochondrial permeability transition pore (mPTP) either directly or indirectly (61, 67, 68). This process disrupts mitochondrial functions, including ATP synthesis and mitochondrial membrane potential, ultimately leading to mitochondrial rupture and release of apoptogenic factors such as cytochrome c. In scenarios, where cell lacks glycolytic ATP sources, this cascade results in mPT-driven necrotic cell death. In contrast, with sufficient ATP levels, early membrane damage is blocked, and apoptogenic factor release initiates ATP-dependent caspase activation and apoptosis (69, 70). We show that BteA-triggered elevation of cytosolic Ca^2+^ precedes loss of mitochondrial membrane potential and early membrane permeabilization. Transmission electron microscopy additionally confirms mitochondrial swelling and cristolysis. These results demonstrate that BteA induces mitochondrial dysfunction, consistent with the concept of mitochondrial Ca^2+^ overload and mPTP-mediated necrosis.

### Limitations of the study

Although our study provides substantial insights into the cytotoxicity of the *Bordetella* type III effector protein BteA, the experiments were primarily carried out using HeLa and HEK cells. While these cell lines are valuable for mechanistic studies and analyzing the disruption of cellular signaling by virulence factors, they do not fully replicate the complexity of an *in vivo* environment. This includes the resistance of certain cell types to Ca^2+^ cytotoxicity and their capacity to maintain ATP levels.

## Methods

### Bacterial strains and growth conditions

The bacterial strains used in this study are listed in Table S2. *Escherichia coli* strain XL1-Blue was used for plasmid construction, and *E. coli* strain SM10λ pir was used for plasmid transfer into *B. bronchiseptica* RB50 by bacterial conjugation. *E. coli* strains were cultivated at 37°C in Luria-Bertani (LB) agar or LB broth. When appropriate, the LB medium was supplemented with chloramphenicol (30 μg/ml), kanamycin (30 µg/ml), or ampicillin (100 µg/ml) for *E. coli* XL1 Blue, and chloramphenicol (15 μg/ml) for *E. coli* SM10λ. *B. bronchiseptica* RB50 and *B. pertussis* B1917 strains were grown on Bordet-Gengou (BG) agar medium (Difco, USA) supplemented with 1% glycerol and 15% defibrinated sheep blood (Lab-MediaServis, Jaromer, Czech Republic) at 37°C and 5% CO_2_. Liquid cultures were done in modified Stainer-Scholte (SSM) medium with reduced L—glutamate (monosodium salt) concentration (11.5 mM, 2.14 g/l) and without FeSO_4_.7H_2_O to maximize the expression of T3SS, as reported previously (25). Culture medium of *B. bronchiseptic*a RB50 harboring pBBRI plasmid (Table S2) was further supplemented with chloramphenicol (30 μg/ml).

### Plasmid construction and introduction into *B. bronchiseptica*

Plasmids used in this study are listed in Table S3, and were constructed using Gibson assembly strategy (71). The GroES promotor (391 nt, NC_002927.3, 1 041 354 – 1 041 744) was amplified from chromosomal DNA of *B. bronchiseptica* RB50, whereas the mNeonGreen coding sequence was amplified from 4xmts-mNeonGreen vector (Addgene item # 98876) using Herculase II Phusion DNA polymerase (Agilent, USA). The pBBRI plasmids encoding mScarlet and mNeonGreen fluorescent proteins were introduced into *B. bronchiseptica* cells by conjugation, using *E. coli* SM10 λpir as plasmid donor strain. The *B. bronchiseptica* conjugants were selected on Bordet-Gengou (BG) blood agar plates supplemented with chloramphenicol (60 μg/ml) and cephalexin (10 μg/ml).

### Cell culture and transfection

HeLa (ATCC CCL-2, human cervical adenocarcinoma), 293T (ATCC CRL-3216, human epithelial kidney cell line) and HEK-Cas9 (293[HEK-293] Cas9, ATCC CRL-1573Cas9, human embryonic kidney cell line constitutively expressing Cas9 from *Streptococcus pyogenes*) were cultivated in Dulbecco’s Modified Eagle Medium (DMEM, Sigma, USA) supplemented with 10% fetal bovine serum (DMEM-10%FBS) at 37°C and 5% CO_2_. Transfections of HeLa cells with plasmids encoding fluorescent markers for ER and mitochondria visualization (Table S3), or Ca^2+^ sensors ER-LAR-GECO and mito-LAR-GECO (Table S3), were performed at 50% cell confluency using PEI MAX transfection reagent (Polysciences). After 24 hours, 5x104 of transfected cells were seeded per coverslip in 24-well plates, or 1x105 per well in 4 well glass-bottom dish (Cellvis) and allowed to adhere overnight before infection.

### Time-lapse live cell imaging for determination of morphological changes and plasma membrane integrity

1x105 of HeLa cells were seeded per well in 4 well glass-bottom dish (Cellvis). The next day, *Bb*WT expressing the fluorescent protein mNeonGreen (*Bb*WT / mNG) or its Δ*bteA* mutant (*Bb*Δ*bteA* / mNG) (Table S2) were added at MOI 10:1 along with propidium iodide (PI, 5 µg/ml). After centrifugation (5 min, 120 g), glass-bottom dish was transferred to a prewarmed stage-top incubation chamber of motorized fluorescence microscope (IX-83, Olympus, Japan) equipped with CCD camera Photometrics prime 95B. Bright-field and fluorescence microscopic images were acquired using a 60x oil immersion objective (PLAPON60XOSC2, NA = 1.4) at a temperature 37°C and 5% CO_2_ during imaging process. CellSens software was used for adjustment, image acquisition and recording sequential images in 16-bit mode at 5 min intervals. Fluorescence was captured using quadruple dichroic mirror DAPI/FITC/TRITC/CY5 with filter block exc. 555/25 nm, em. 605/52 nm for PI, and exc. 490/20 nm, em. 525/36 nm for mNG.

### Ninety-six well plate cytotoxicity assay

Cytotoxicity of *B. bronchiseptica* and *B. pertussis* towards HeLa and HEK-Cas9 was determined as changes in cell membrane integrity using the fluorescent DNA binding dye CellTox Green (Promega, Cat. No. G8743), as described previously (29). In brief, 2 x 104 of HeLa or HEK-Cas9 cells per well were seeded in a 96-well black/clear bottom plate (Corning, USA) in DMEM-10%FBS. The next day, *B. bronchiseptica* or *B. pertussis* strains were added at the indicated MOI along with CellTox Green reagent. The plate was centrifuged (5 min, 120 g) and placed inside the chamber with 37°C and 5% CO_2_ of the TecanSpark microplate reader (Tecan, Switzerland). Fluorescence measurements at 494ex/516em with a 10 nm bandwidth for both were performed at defined time intervals for 10 h. If appropriate, HeLa cells were pre-treated with 2-aminoethyl diphenylborinate (2-APB) at 100 µM for 30 min, or ruthenium 360 at 10 µM, and cyclosporin A at 1 µM for 60 min prior to *B. bronchiseptica* infection. Additionally, glycine at 5 mM was added 5 min before infection when required.

### Visualisation of cellular organelles in fixed cells

5x104 of HeLa cells transiently expressing fluorescent markers for ER and mitochondria visualization (Table S3) were seeded per coverslip and incubated at 37°C and 5% CO_2_. The next day, HeLa cells were infected with the indicated *B. bronchiseptica* or *B. pertussis* strains at a MOI 50:1. Added bacteria were centrifugated (5 min, 120 g) onto the cell surface, and samples were fixed by 4 % formaldehyde solution in PBS (20 min, RT) 1 and 5 h post-infection. Coverslips were then rinsed with distilled water and mounted onto microscope glass slides using Vectashield mounting medium (Vector Laboratories, USA). Fluorescence microscopy was performed using a motorized fluorescence microscope (IX-83, Olympus, Japan) equipped with CCD camera Photometrics prime 95B with 100x oil-immersion objective (UPLXAPO100XOPH, N.A. = 1.45). Fluorescence was captured using quadruple dichroic mirror DAPI/FITC/TRITC/CY5 with filter block exc. 555/25 nm, em. 605/52 nm for mScarlet, and exc. 490/20 nm, em. 525/36 nm for mNG. Images were collected in a 16-bit format using CellSens software, with Z-stacks taken with 0.26 µm z-steps, and deconvolution performed with Advanced Maximum Likelihood (AMLE) filters. A single focal plane of a Z-stack is presented in all figures.

### Production of lentiviral vector library and genome-wide CRISPR-Cas9 knockout screen in HEK-Cas9 cells

The production of lentiviral vector library GeCKO v2 (Addgene item #1000000049, (35)) consisting of sublibraries A and B and preparation of the corresponding sublibraries of HEK-Cas9 cells were done according to the protocol reported by Joung *et al*. (72). In brief, 6x106 of 293T cells per T75 flask were seeded, with total of twenty-four T75 flask utilized. The next day, each of the T75 flask was co-transfected with 4.5 µg of pCMV-VSV-G, 6.5 µg of psPAX2 and 9 µg of either GeCKO v2 human sublibrary A or B (Table S3) using PEI MAX transfection reagent (Polysciences). After 48 hours, the supernatant of the 293T cells containing packaged vector sublibrary A or B was collected, filtered through a 0.45 µm filter, and stored at -80°C. To determine the viral titer in the collected supernatants, HEK-Cas9 cells were transduced in a 96-well white/clear bottom plate (Corning, USA) using varying amounts of the supernatant in the presence of polybrene (8 μg/ml). Following supernatant addition, cells were centrifuged (2 h, 1000 g) and cultivated for 24 hours before selection with puromycin (0.6 µg/ml). After 5 days, the number of surviving cells was assessed using Cell Titer Glo (Promega, Cat.no. G7570), and virus titer was calculated.

Subsequently, HEK-Cas9 sublibraries A and B were prepared through lentivirus spin infection of HEK-Cas9 cells at MOI of 0.3:1 to ensure a single sgRNA delivery per cell, as follows. HEK-Cas9 cells were seeded into 6-well plates, lentivirus was added in the presence of polybrene (8 μg/ml), and plates were centrifugated (2 h, 1000 g). A total of 108 of HEK-Cas9 cells were transduced per sublibrary to achieve coverage of >500 cells per sgRNA. Twenty-four hours post-transduction, HEK-Cas9 cells were selected with puromycin (0.6 µg/ml) for 5 days. The surviving cells were expanded and HEK-Cas9 libraries were cryostocked at 108 of cells per sublibrary stock. The preservation of complexity in the sublibraries was next assessed by next-generation sequencing. In brief, genomic DNA from 108 cells per sublibrary was harvested using Blood & Cell culture DNA Maxi kit (QIAGEN, Cat.no. 13362), and the sgRNA regions were amplified and barcoded using a PCR reaction, as described in (72). The barcoded PCR products were pooled in equimolar ratios and concentrated using Zymo-Spin V Columns (Zymo Research, Cat.no. C1012-25). The concentrated PCR products were separated on a 2% agarose gel and extracted using the DF300 Gel/PCR DNA Fragments Extraction Kit (Geneaid, Cat.no. DF300). DNA concentration in the samples was determined by Qubit dsDNA HS Assay Kit (Invitrogen), and product length was verified using Agilent High Sensitivity DNA Kit. The samples were sequenced on the Illumina NextSeq 2000 platform (single-end, 100 cycles). Sequence data were analyzed and aligned to the sequence reads in a reference file of all sgRNA sequences in sublibrary A and B, which is provided by Addgene.

For positive selection of prepared GeCKO v2 HEK-Cas9 sublibraries A and B by infection with *B. brochiseptica* RB50 wild type, we employed 108 of cells per HEK-Cas9 sublibrary (ensuring > 1,500 coverage per sgRNA). Each HEK-Cas9 sublibrary was distributed across five 15 cm Petri dishes and infected with *Bb*WT at MOI of 100:1. The infection was allowed to proceed at 37°C and 5% CO_2_ for 5 h. To stop the infection, the medium containing *Bb*WT was discarded and replaced with fresh medium containing gentamicin (100 µg/ml). These conditions resulted in the death of approximately 95% of infected cells. Surviving cells were expanded to 108 cells per sublibrary and subjected to another round of infection with *Bb*WT. This process was repeated five times, after which the sensitivity of selected HEK-Cas9 sublibrary cells was assessed using lactate dehydrogenase (LDH) assay. For LDH assay, parental and selected HEK-Cas9 sublibrary A and B cells were seeded at 106 per well in 6-well plate in DMEM with 2% (vol/vol) FBS without phenol red indicator. The next day, *Bb* strains were added at indicated MOI, and infection was allowed to proceed at 37°C and 5% CO_2_ for 6h. The LDH release into cell culture media was determined using CytoTox 96 assay (Cat.No. G1780, Promega) according to the manufacturer instructions. The % cytotoxicity was calculated using the following equation: (OD495 sample – OD495 media)/(OD495 total lysis – OD495 media)* 100.

### Cytosolic Fluo-4/AM calcium assay

Cytosolic calcium influx induced by *B. bronchiseptica* infection was assessed through fluorescence intensity measurement and time-lapse imaging using the Fluo-4 Direct Calcium Assay (Cat. No. F10471, Invitrogen), according to manufacturer instructions. In brief, 2 x 104 of HeLa cells in DMEM-10%FBS were seeded per well in a 96-well black/clear bottom plate (Corning, USA) or 1 x 104 of cells per well in 4-well glass-bottom dish (Cellvis). The next day, cells were loaded with Fluo-4 Direct calcium loading solution supplemented with 5 mM probenecid for 1 h at 37°C and 5% CO_2_. *B. bronchiseptica* strains were then added at the indicated MOI in Fluo-4 Direct calcium reagent and either centrifuged onto the HeLa cell surface (5 min, 120 g) or added directly without centrifugation, as indicated. For fluorescence-intensity measurement, 96-well plate was placed inside the chamber with 37°C and 5% CO_2_ of the TecanSpark microplate reader (Tecan, Switzerland), and fluorescence measurements at 494ex/516em with a 10 nm bandwidth for both were taken at 2-min intervals over a period of 10 h. If appropriate, HeLa cells were pre-treated with 2-aminoethyl diphenylborinate (2-APB) at 100 µM for 30 min prior to *B. bronchiseptica* infection. Additionally, glycine at 5 mM was added 5 min before infection when required. For time-lapse imaging, fluorescent signals were captured using a motorized fluorescence microscope (IX-83, Olympus, Japan) equipped with CCD camera Photometrics prime 95B with 30x silicon immersion objective (UPLSAPO30XS, NA = 1.05) at a temperature 37°C and 5% CO_2_ during imaging process. CellSens software was used for adjustment, image acquisition and recording sequential images in 16-bit mode at 2 min time intervals. Fluorescence was captured using quadruple dichroic mirror DAPI/FITC/TRITC/CY5 with filter block exc. 555/25 nm, em. 605/52 nm for mScarlet-expressing bacteria, and exc. 490/20 nm, em. 525/36 nm for Fluo-4/AM.

### Visualization of calcium ion fluxes, mitochondrial morphology and mitochondrial membrane potential

To examine calcium ion fluxes, a plasmid-encoded red fluorescent Ca^2+^ sensor targeted to either the ER (ER-LAR-Geco) or mitochondria (mito-LAR-Geco) (Table S3) was used in combination with cytosolic Fluo-4/AM calcium assay. Sensor-transfected HeLa cells were seeded on 4 well glass-bottom dish (Cellvis) and loaded with Fluo-4 Direct calcium loading solution supplemented with 5 mM probenecid for 1 h. Infection with *B. bronchiseptica* was performed at a MOI 10:1. Alternatively, mitochondrial morphology or membrane potential was assessed by staining with MitoTracker Orange CM-H2TMRos (MitoTracker, 500 nM, Invitrogen, Cat. No. M7511) or Tetramethylrhodamine (TMRM, 100 nM, Invitrogen, Cat. No. T668) according to the manufacturer instructions in combination with Fluo-4/AM staining. Fluorescence signals were captured using a motorized fluorescence microscope (IX-83, Olympus, Japan) as described above using 40x dry objective (UPLXAPO40X, NA = 0.95) or 60x oil objective (PLAPON60XOSC2, NA = 1.4) at a temperature 37°C and 5% CO2 during imaging process. Fluorescence was captured with filter block exc. 555/25 nm, em. 605/52 nm for ER-LAR-Geco, mito-LAR-Geco, MitoTracker, and TMRM, and exc. 490/20 nm, em. 525/36 nm for Fluo-4/AM. Images were collected in a 16-bit format using CellSens software at 3 min time intervals.

### Transmission electron microscopy

5x104 of HeLa cells were seeded per coverslip in 24-well plates and incubated at 37°C and 5% CO_2_. The next day, cells were infected with *Bb*WT and Δ*bteA* stains at MOI 10:1, or left untreated. Added bacteria were centrifugated (5 min, 120 g) onto the cell surface, and samples were fixed 1 h post-infection by 2% glutaraldehyde, 4% paraformaldehyde in 2 mM CaCl_2_, 100 mM Hepes pH 7.4 for 2 h. After washing with bi-distilled water, the cells were postfixed in 1% osmium tetroxide, dehydrated with a series of acetone and embedded in Epon-Durcupan resin (Sigma-Aldrich, St. Louis, MO, USA). The resin was heat-polymerized at 60°C for 72 hours. Ultrathin sections (90 nm) were prepared using a Leica EM UC6 ultramicrotome (Leica Microsystems, Wetzlar, Germany) with a diamond knife (Diatome, Biel, Switzerland). The sections were mounted on 200 mesh size copper grids and examined in a JEOL JEM-1400 Flash transmission electron microscope operated at 80 kV, equipped with a Matataki Flash sCMOS camera (JEOL, Akishima, Tokyo, Japan).

### Image processing

All image processing steps were performed using Fiji (73).

## Acknowledgments

This work was supported by the grant 21-05466S of the Czech Science Foundation (www.gacr.cz), grant Talking microbes - understanding microbial interactions within One Health framework (CZ.02.01.01/00/22_008/0004597) of the Ministry of Education, Youth and Sports of the Czech Republic (www.msmt.cz) and the Lumina Queruntur Fellowship LQ200202001 of the Czech Academy of Sciences to J.K. O.C. was funded by ESF International Mobility of Researchers, grant number CZ.02.2.69/0.0/0.0/18_053/0017705. We would also like to acknowledge the support of the project LM2023053 (Czech National Node to the European Infrastructure for Translational Medicine) from Ministry of Education, Youth and Sports of the Czech Republic, and the Electron Microscopy Core Facility, IMG ASCR, Prague, CR, supported by MEYS CR (LM2018129, LM2023050, CZ.02.1.01/0.0/0.0/18_046/0016045, CZ.02.1.01/0.0/0.0/16_013/0001775), particularly the contributions of Dominik Pinkas. The funders had no role in study design, data collection and analysis, decision to publish, or preparation of the manuscript.

## Author contributions

Conceptualization: JK; Data curation: MZ, JK; Formal Analysis: MZ, ES, BP, JK; Funding acquisition: JK; Investigation: MZ, ES, BP, MC, OC, TRA, JK; Methodology: OC, TG, IM, JK; Project administration: JK; Resources: JK; Software: MZ; Supervision: OC, TG, JK; Validation: MZ, ES, TG, JK; Visualization: MZ, JK; Writing: JK; Writing – review & editing: MZ, OC.

## Inclusion and diversity statement

Figures were generated using a color-blind accessible palette, promoting inclusion and diversity in data visualization.

---

**Suppl. Fig. 1.**
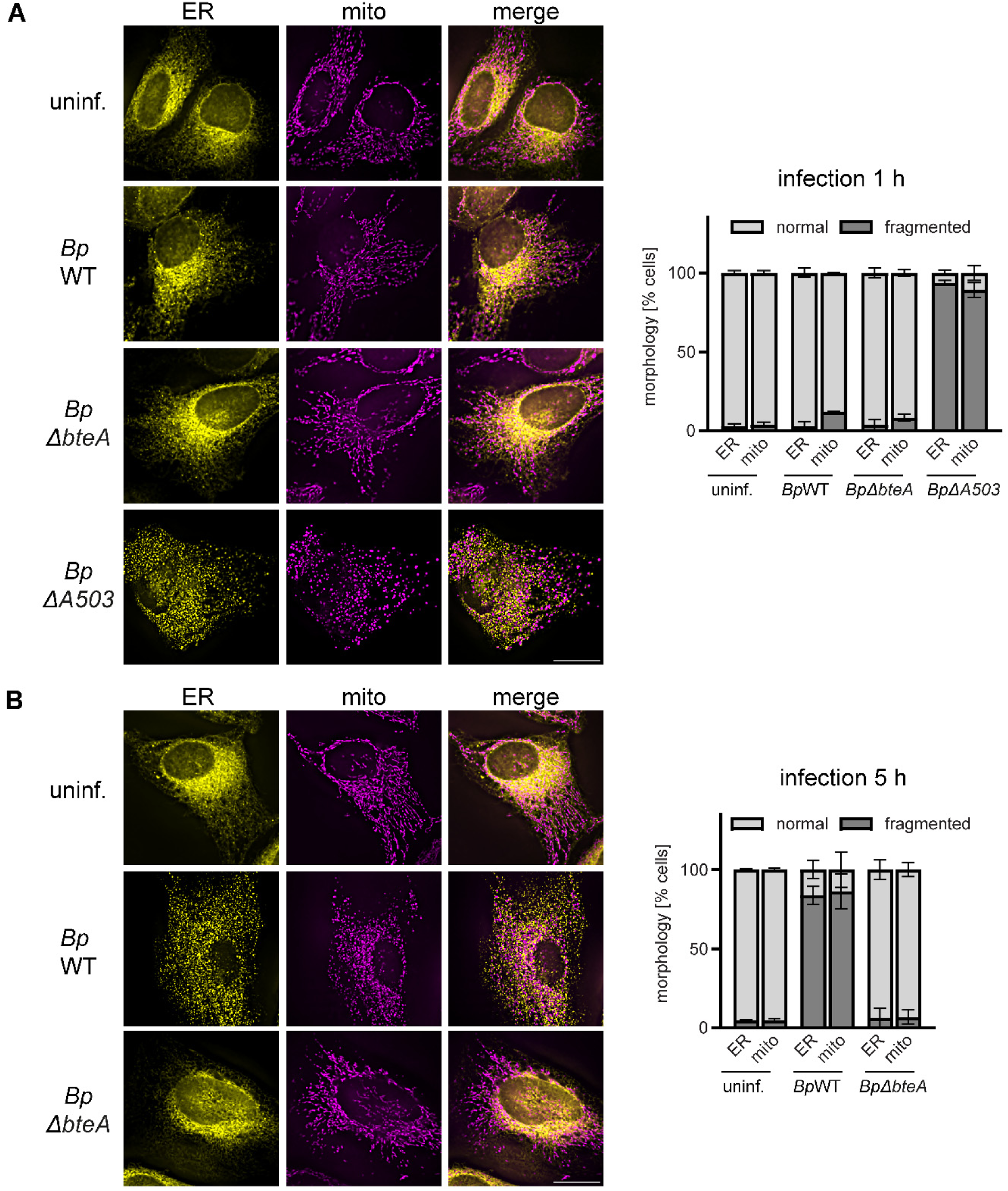
BteA-induced cell death is characterized by fragmentation of endoplasmatic reticulum and mitochondrial networks. Hela cells were transfected to express fluorescent proteins tagged with localization signals for endoplasmatic reticulum (ER) and mitochondria (mito). Cells were infected with the indicated *Bp* strains at an MOI 50:1 for 1 h **(A)** or 5 h **(B)**, or left untreated, followed by fixation and analysis by fluorescence imaging. The shown micrographs are representative of two independent experiments from which the organelle morphology was determined. Data represent the mean ± SEM of at least 100 cells counted per experiment. ER, yellow; mitrochondria, magenta. Scale bar, 20 µm.

**Suppl. Fig. 2.**
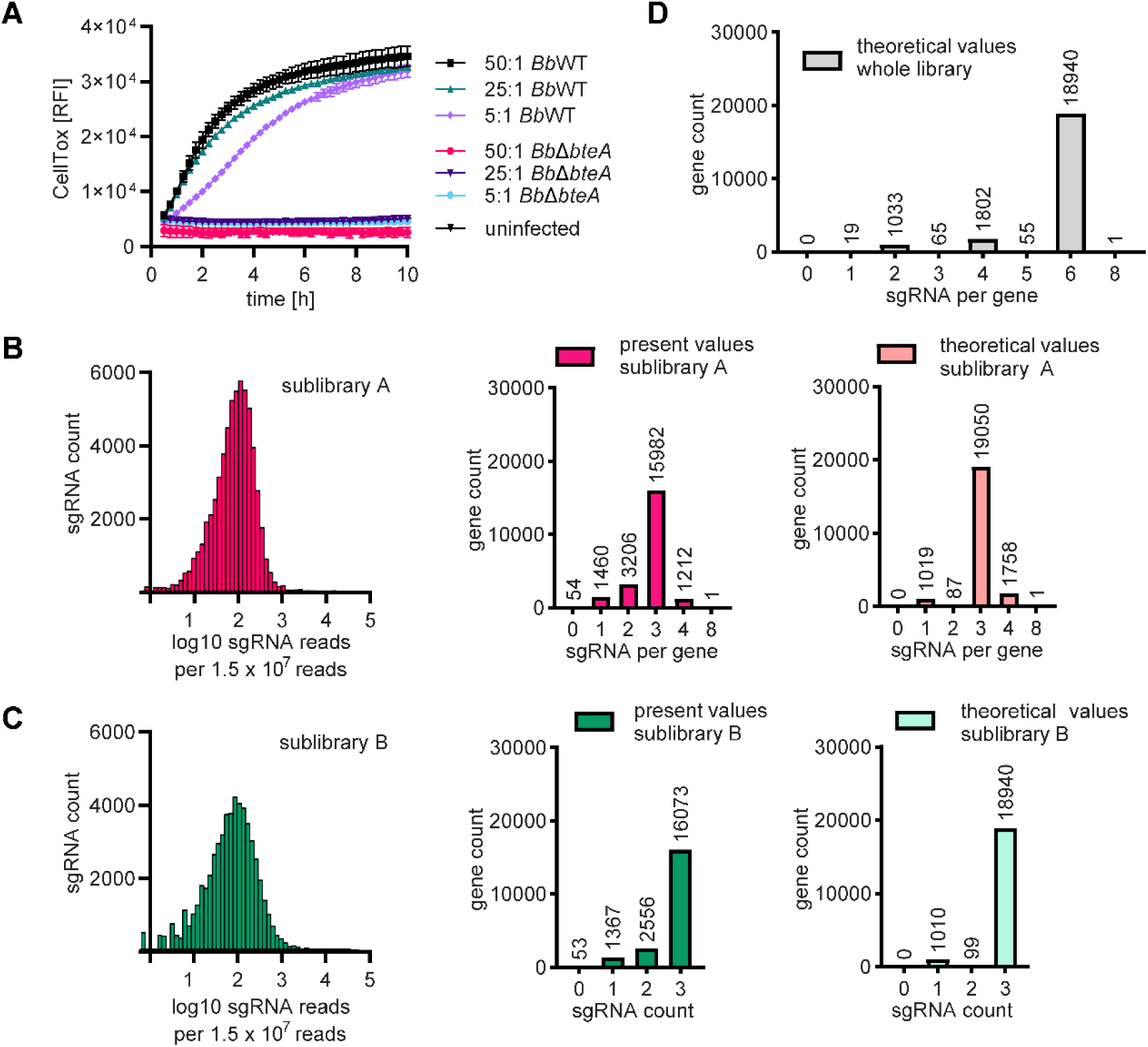
Susceptibility of Hek-Cas9 cells to BbWT infection and characterization of Hek-Cas9 library complexity. **(A)** Susceptibility of Hek-Cas9 cells. Hek-Cas9 cells were infected with *B. bronchiseptica* strains at the indicated MOI. Plasma membrane permeabilization was determined using the fluorescent DNA binding dye CellTox Green. Data represent the mean ± SEM of a representative experiment from 2 independent experiments. **(B-C)** Complexity in the individual Hek-Cas9 sublibraries A and B. Distribution of detected sgRNA is depicted in the histograms on the left. These histograms show the number of individual sgRNAs against their detection frequency. Non-detected sgRNA are shown as well. The middle graphs display the number of targeted genes per detected sgRNA count, and these are compared to the theoretically expected values shown in the right graphs. **(D)** Theoretical complexity of the combined sublibraries A and B. The number of targeted genes per theoretical sgRNA count is indicated. Please compare with Fig. 2B in the main text.

**Suppl. Figure 3.**
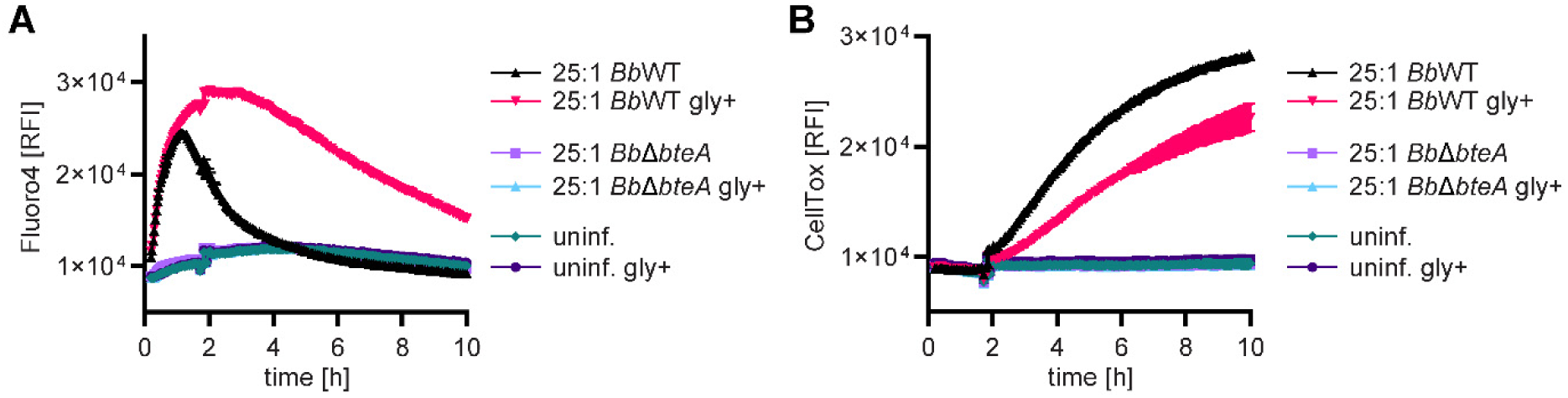
Inhibition of cell plasma membrane permeabilization by glycine does not diminish calcium influx. HeLa cells were infected with *Bb*WT or *Bb*Δ*bteA* derivative at MOI 25:1 in the presence (gly+) or absence of 5 mM glycine. Calcium influx was assessed using Fluo-4/AM Ca^2+^ indicator **(A)** whereas plasma membrane permeabilization was determined in parallel wells by fluorescent DNA binding dye CellTox Green **(B)**. Data represent the mean ± SEM of a representative experiment from 2 independent experiments.

**Suppl. Fig. 4.**
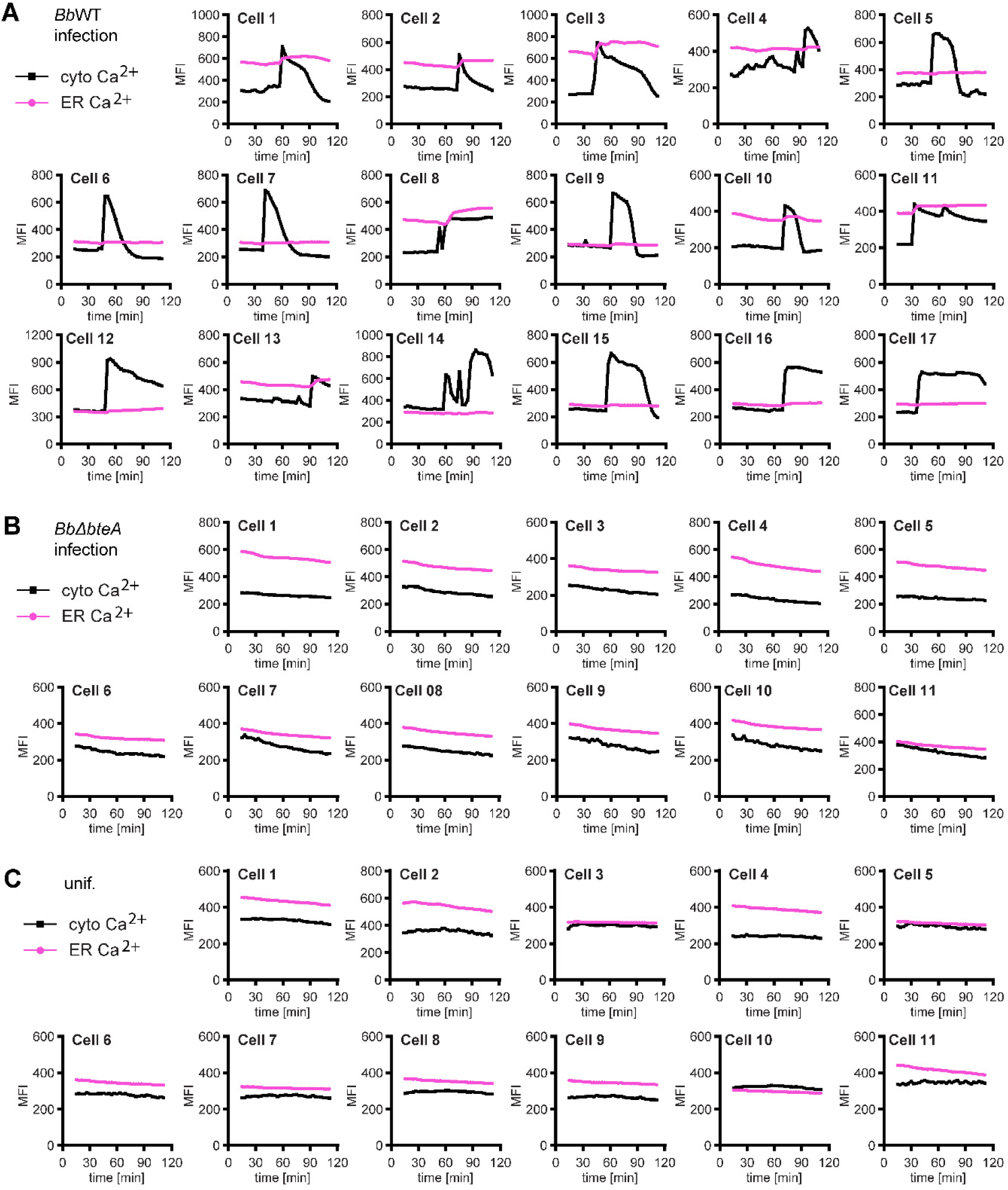
Correlation of cytosolic and ER calcium levels in individual cells. Hela cells, transfected to express ER-targeted red Ca^2+^ sensor ER-LAR-GECO, were loaded with the cytosolic Ca²⁺ indicator Fluo-4/AM, and infected with *Bb*WT **(A)** or *BbΔbteA* **(B)** strains at MOI of 10:1, or left untreated **(C)**. The graphs indicate the mean fluorescence intensities of the individual cells quantified over time.

**Suppl. Fig. 5.**
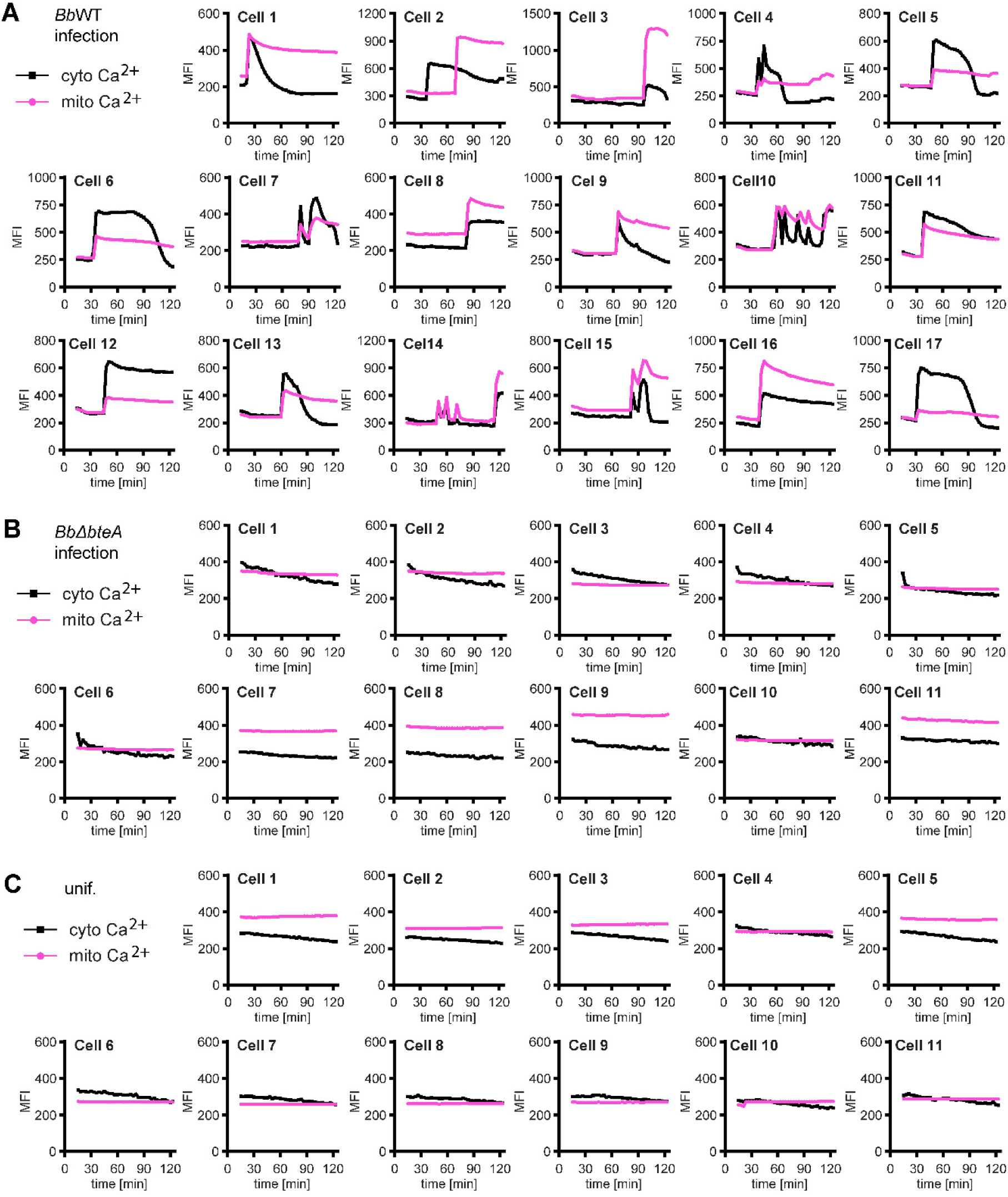
Correlation of cytosolic and mitochondrial calcium levels in individual cells. Hela cells, transfected to express mitochondria-targeted red Ca^2+^ sensor mito-LAR-GECO, were loaded with the cytosolic Ca²⁺ indicator Fluo-4/AM, and infected with *Bb*WT **(A)** or *BbΔbteA* **(B)** strains at MOI of 10:1, or left untreated **(C)**. The graphs indicate the mean fluorescence intensities of the individual cells quantified over time.

**Suppl. Fig. 6.**
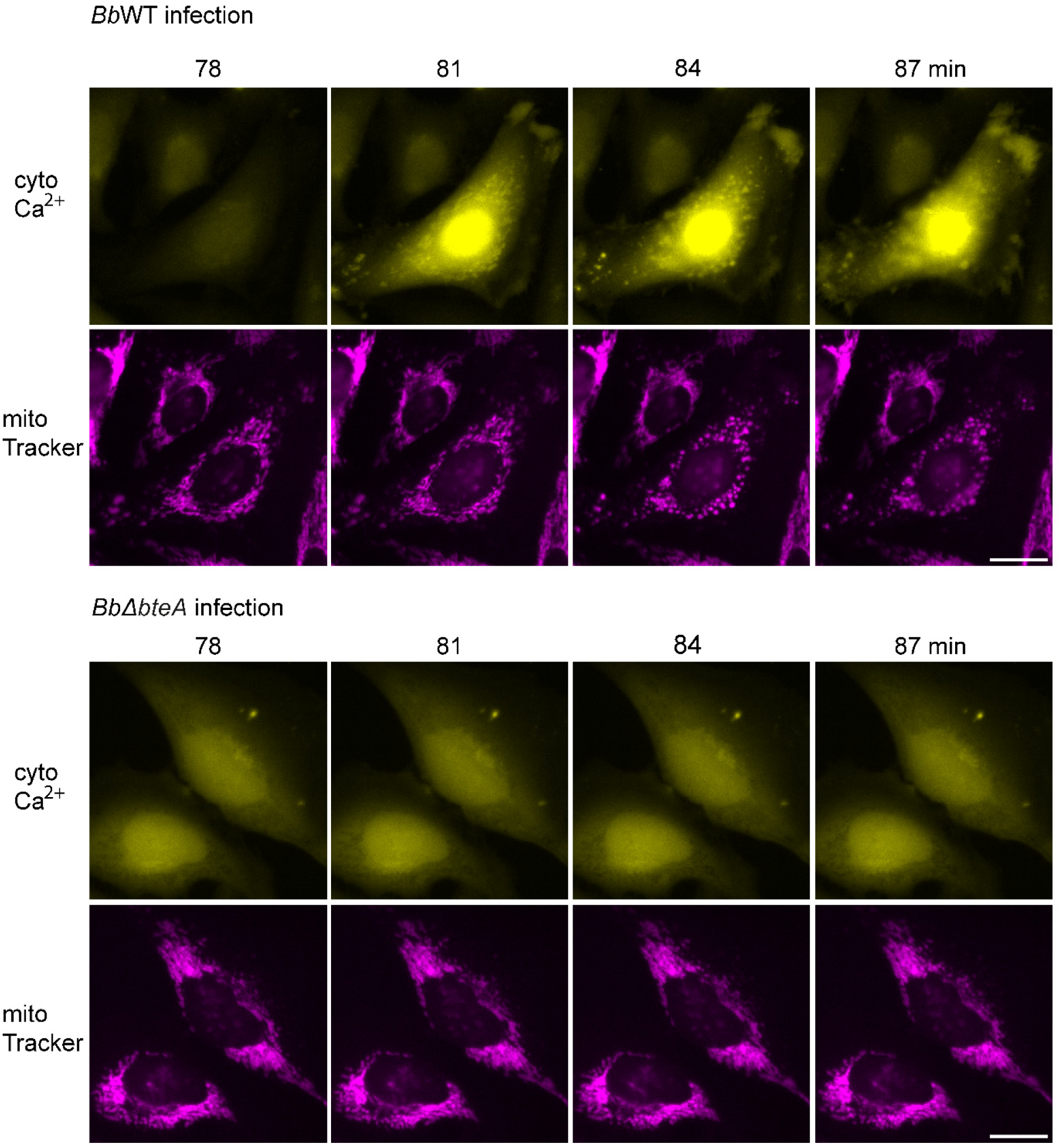
Assessment of mitochondrial morphology during *Bb* infection. HeLa cells were loaded with MitoTracker, and the cytosolic Ca^2+^ indicator Fluo-4/AM, before being infected with *Bb*WT or *Bb*Δ*bteA* derivative at MOI 10:1. Sequence of time lapse images is shown. Data are representative of two independent experiments. Cytosolic Ca^2+^ indicator Fluo-4/AM, yellow; MitoTracker, magenta. Scale bar, 20 µm.

**Suppl. Fig. 7.**
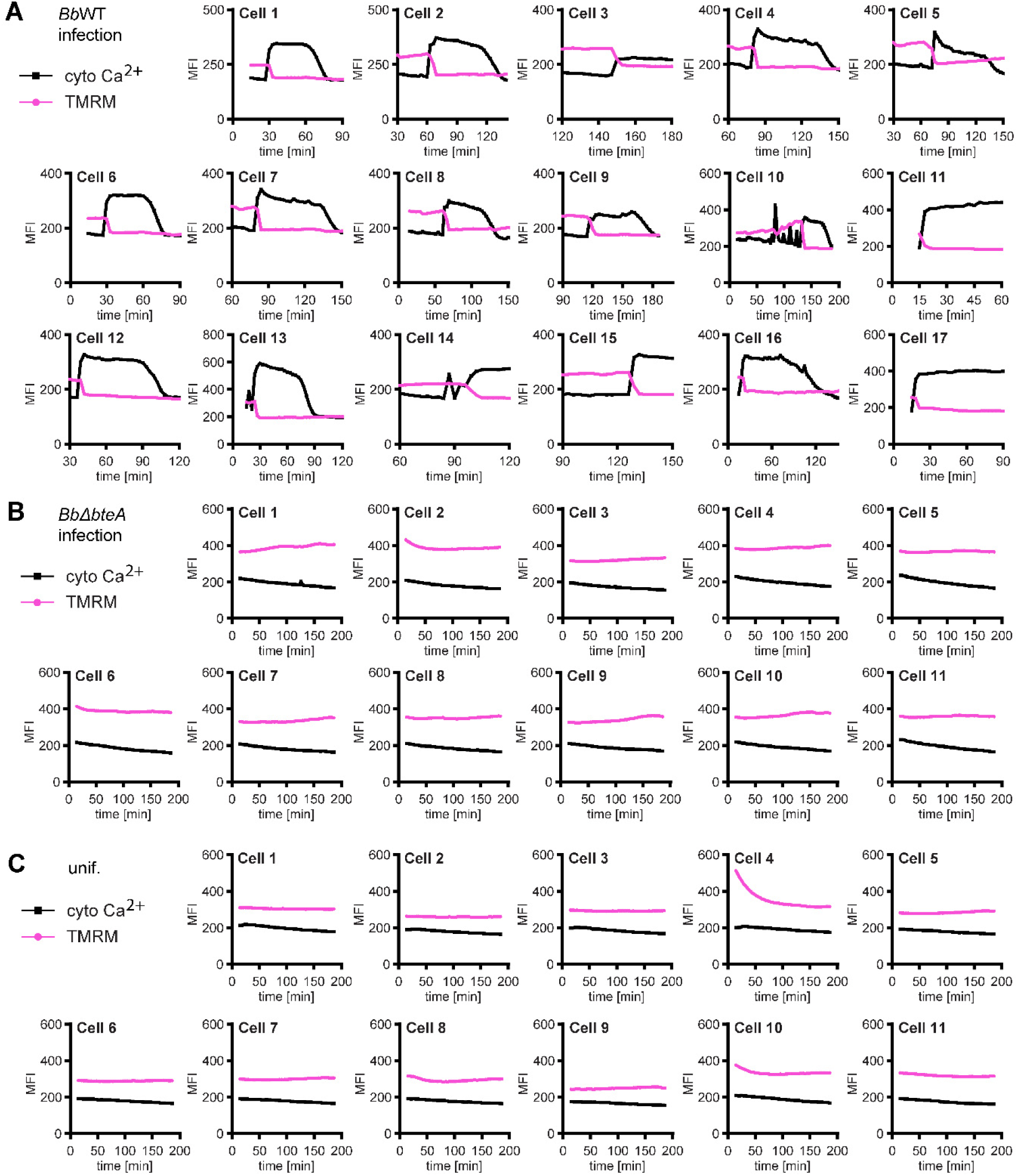
Correlation of cytosolic calcium levels and mitochondrial membrane potential in individual cells. HeLa cells were loaded with the mitochondrial membrane potential indicator TMRM and the cytosolic Ca^2+^ indicator Fluo-4/AM, and infected with *Bb*WT **(A)** or *BbΔbteA* **(B)** strains at MOI of 10:1, or left untreated **(C)**. The graphs indicate the mean fluorescence intensities of the individual cells quantified over time.

**Suppl. Figure 8.**
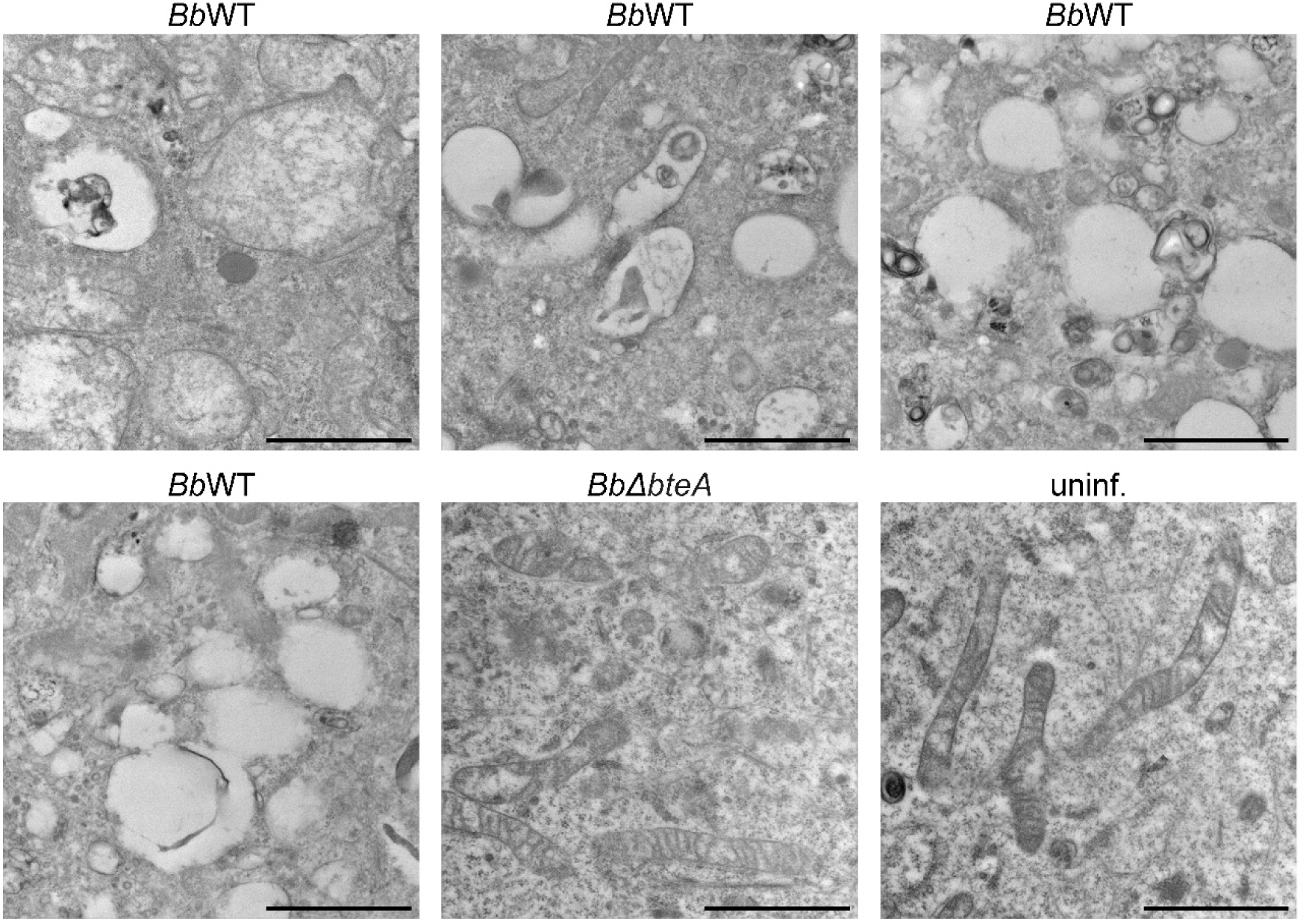
Transmission electron microscopy images of mitochondria. Hela cells were infected with *Bb*WT and *Bb*Δ*bteA* at MOI of 25:1 for 1 h or left untreated. Following fixation, ultrathin sections were prepared and analyzed. The swelling of mitochondria with cristolysis induced by BteA was observed. Images are representative of two independent experiments. Scale bar, 1 µm.

## List of Supplementary Videos

**Suppl. Video 1. Imaging of morphological changes and plasma membrane permeability in HeLa cells.** HeLa cells were infected with *B. bronchiseptica* wild type (*Bb*WT) and *Bb*Δ*bteA* mutant (*Bb*Δ*bteA*), expressing the fluorescent protein mNeonGreen, at MOI of 10:1 in the presence of propidium iodide (5 µg/ml). Bacteria were centrifugated onto cell surface. Images were captured every 5 min and are shown at 3 frames per second. Video is representative of three independent experiments. Bright field, gray; bacteria, cyan; propidium iodide, magenta. Scale bar, 20 µm.

**Suppl. Video 2. Calcium imaging.** HeLa cells, loaded with cytosolic Ca^2+^ indicator Fluo-4/AM, were infected with *Bb*WT and *Bb*Δ*bteA,* which express the fluorescent protein mScarlet, at MOI of 10:1. To allow tracking bacterial attachment, no centrifugation of bacteria onto the cell surface was performed. Images were captured at 2 min intervals and are displayed at 3 frames per second. This video is representative of two independent experiments. Bacteria, magenta; cytosolic Ca^2+^ indicator Fluo-4/AM, yellow. Scale bar, 20 µm.

**Suppl. Video 3. Calcium imaging.** HeLa cells, loaded with cytosolic Ca^2+^ indicator Fluo-4/AM, were infected with mScarlet-expressing *Bb*WT and *Bb*Δ*bteA* at MOI of 10:1. Bacteria were centrifugated onto cell surface. Images were captured every 2 min and are shown at 3 frames per second. Video is representative of three independent experiments. Bacteria, magenta; cytosolic Ca^2+^ indicator Fluo-4/AM, yellow. Scale bar, 20 µm.

**Suppl. Video 4. ER calcium imaging.** HeLa cells, transfected to express red fluorescent Ca^2+^ sensor targeted to ER (ER-LAR-Geco), were loaded with cytosolic Ca^2+^ indicator Fluo-4/AM. Infection with *Bb*WT and *Bb*Δ*bteA* was performed at MOI 10:1, followed by bacterial centrifugation onto cell surface. Images were captured at 3 min intervals and are displayed at 3 frames per second. Video is representative of two independent experiments. Cytosolic Ca^2+^ indicator Fluo-4/AM, yellow; ER Ca^2+^ sensor, magenta. Scale bar, 20 µm.

**Suppl. Video 5. Mitochondrial calcium imaging.** HeLa cells, transfected to express red fluorescent Ca^2+^ sensor targeted to mitochondria (mito-LAR-Geco), were loaded with cytosolic Ca^2+^ indicator Fluo-4/AM. Infection with *Bb*WT and *Bb*Δ*bteA* was performed at MOI 10:1, followed by bacterial centrifugation onto cell surface. Images were captured at 3 min intervals and are displayed at 3 frames per second. This video is representative of two independent experiments. Cytosolic Ca^2+^ indicator Fluo-4/AM, yellow; mitochondria Ca^2+^ sensor, magenta. Scale bar, 20 µm.

**Suppl. Video 6. Imaging of mitochondria morphology.** HeLa cells were loaded with the MitoTracker and the cytosolic Ca*2+* indicator Fluo-4/AM, before being infected with *Bb*WT and *Bb*Δ*bteA* derivative at MOI 10:1. Bacteria were centrifugated onto cell surface. Images were captured at 3 min intervals and are displayed at 3 frames per second. Video is representative of three independent experiments. MitoTracker, magenta; cytosolic Ca^2+^ indicator Fluo-4/AM, yellow. Scale bar, 20 µm.

**Suppl. Video 7. Imaging of mitochondrial membrane potential.** HeLa cells were loaded with the mitochondrial membrane potential probe TMRM, and the cytosolic Ca*2+* indicator Fluo-4/AM, before being infected with *Bb*WT and *Bb*Δ*bteA* derivative at MOI 10:1. Bacteria were centrifugated onto cell surface. Images were captured at 3 min intervals and are displayed at 3 frames per second. Video is representative of three independent experiments. TMRM, magenta; cytosolic Ca^2+^ indicator Fluo-4/AM, yellow. Scale bar, 20 µm.

**Suppl. Video 8. Imaging of mitochondrial membrane potential.** HeLa cells were pre-incubated with the inhibitor of mitochondrial calcium uniporter ruthenium 360 (Ru360+) at 10 µM or left untreated. Subsequently, the cells were loaded with TMRM, and the cytosolic Ca*2+* indicator Fluo-4/AM, before being infected with *Bb*WT at MOI 10:1. Bacteria were centrifugated onto cell surface. Images were captured at 3 min intervals and are displayed at 3 frames per second. The video is representative of two independent experiments. TMRM, magenta; cytosolic Ca^2+^ indicator Fluo-4/AM, yellow. Scale bar, 20 µm.

**S1 Table.** List of genes for which no sgRNA was detected in the combined sublibraries A and B despite their theoretical presence.

| Gene_ID | Gene_name | Description |
| --- | --- | --- |
| PLN | Phospholamban | Inhibitor of cardiac sarcoplasmic reticulum Ca<sup>2+</sup> pump (1) |
| hsa-mir-1207 | MicroRNA 1207 | unknown |
| hsa-mir-1299 | MicroRNA 1299 | regulation of tumor pathogenesis, tumor suppressor (2) |
| hsa-mir-4679-1 | MicroRNA 4679-1 | unknown |
| hsa-mir-4679-2 | MicroRNA 4679-2 | unknown |
| hsa-mir-5096 | MicroRNA 5096 | unknown |
| hsa-mir-5591 | MicroRNA 5591 | unknown |
| hsa-mir-6088 | MicroRNA 6088 | unknown |
| hsa-mir-8078 | MicroRNA 8078 | unknown |
| NonTargetingControlGuide<br>ForHuman_0583 | - | Non-targeting control guide |

**S2 Table.** List of bacterial strains used in this study.

| Strain | Genotype and relevant description | Internal number | Reference |
| --- | --- | --- | --- |
| ***E. coli* strains** |  |  |  |
| XL1-Blue | *recA1 endA1 gyrA96 thi-1 hsdR17 supE44 relA1 lac F' proAB lacIqZΔM15 Tn10 Tet<sup>r</sup>* |  | Stratagene |
| SM10 λpir | *thi thr leu tonA lacY supE recA::RP4-2-Tc::Mu Km λpir* |  | (3, 4) |
| ***B. bronchiseptica* strains** |  |  |  |
| *BbWT* | *BbRB50* WT; wild type *Bordetella bronchiseptica* RB50 (B1976); complex I rabbit isolate; ST-12 | BB012 | (5, 6) |
| *BbΔbteA* | *BbRB50 ΔbteA*; *BbRB50* strain derivative with *bteA* in-frame deletion of codons L2-A657 | BB020 | (7) |
| *BbWT* / mNG | *BbRB50* WT harboring pBBRI-encoded mNeonGreen (mNG) fluorescent protein under the control of *BbRB50* GroES promoter (PgroES) | pBB119 | This study |
| *BbΔbteA* / mNG | *BbRB50 ΔbteA* harboring pBBRI-encoded mNeonGreen (mNG) fluorescent protein under the control of *BbRB50* groES promoter (PgroES) | pBB121 | This study |
| *BbWT* / mSc | *BbRB50* WT harboring pBBRI-encoded mScarlet (mSc) under the control of *BpTohamal* BvgAS-regulated filamentous hemagglutinin (*fhaB* gene) promoter (PfhaB) | pBB039 | This study |
| *BbΔbteA* / mSc | *BbRB50 ΔbteA* derivative harboring pBBRI-encoded mScarlet (mSc) under the control of *BpTohamal* BvgAS-regulated filamentous hemagglutinin (*fhaB* gene) promoter (PfhaB) | pBB117 | This study |
| ***Bordetella pertussis* strains** |  |  |  |
| *Bp*WT | *BpB1917* WT; wild type *Bordetella pertussis* 1917; *fim2-1*, *fim3-2*, *ptxP3*, *ptxA1*, *ptxB2*, *ptxC2*, *ptxD1*, *ptxE1*, *prn2* | BP001 | (8, 9) |
| *BpΔbteA* | *BpB1917 ΔbteA*; *BpB1917* strain derivative with *bteA* in-frame deletion of codons L2-A656 | BP003 | (10) |
| *BpΔA503* | *BpB1917 bteA ΔA503*; *BpB1917* strain derivative with *bteA* in-frame deletion of codon A503 | BP006 | (10) |

**S3 Table.** List of plasmids used in this study.

| Plasmid | Description | Reference |
| --- | --- | --- |
| pBBRI MCS | *lacPOZ'* *mob*<sup>+</sup>, broad-host cloning vector, Cm<sup>R</sup> | (11, 12) |
| pBBRI-PgroES-mNG | pBBRI vector with *BbRB50* promoter GroES (PgroES) and coding sequence of the mNeonGreen protein (mNG) | This study |
| pBBRI-PfhaB-mSc | pBBRI vector with *BpTohamal* filamentous hemagglutinin promoter (pfhaB) and coding sequence of the mScarlet protein (mSc) | (13) |
| export&KDEL-mScarlet | ER-mScarlet1, pEGFP-N1-derived vector encoding mScarlet fluorescent protein targeted to mitochondria, Addgene item # 137805 | (14) |
| 4xmts-mNeonGreen | 4xmts-mNeonGreen, pEGFP-N1-derived vector encoding mNeonGreen fluorescent protein targeted to mitochondria, Addgene item # 98876 | (14) |
| GeCKO v2 sublibrary A | Human CRISPR Knockout Pooled Library (GeCKO v2) in lentiGuide-Puro A, Catalog #1000000049, Addgene item #52959 | (15) |
| GeCKO v2 sublibrary B | Human CRISPR Knockout Pooled Library (GeCKO v2) in lentiGuide-Puro B, Catalog #1000000049, Addgene item #52960 | (15) |
| pCMV-VSV-G | Vector encoding envelope protein for producing lentiviral and MuLV retroviral particles, Addgene item #8454 | (16) |
| psPAX2 | 2nd generation lentiviral packaging plasmid, Addgene item #12260 | Addgene, Trono Lab unpublished |
| ER-LAR-GECO | CMV-ER-LAR-GECO1, low affinity red intensimetric genetically encoded Ca<sup>2+</sup> indicator targeted to ER, Addgene item #61244 | (17) |
| mito-LAR-GECO | CMV-mito-LAR-GECO1.2, low affinity red intensimetric genetically encoded Ca<sup>2+</sup> indicator targeted to mitochondria, Addgene item #61245 | (17) |

